# Chain-length regulation by WzzE is necessary for, but genetically separable from, cyclic enterobacterial common antigen synthesis

**DOI:** 10.1101/2025.06.25.661564

**Authors:** Joseph F. Carr, Jennifer S. Rudolf, Daniel J. Warzecha, Yohannes H. Rezenom, Angela M. Mitchell

**Affiliations:** Department of Biology, Texas A&M University, College Station, Texas, USA; Chemistry Mass Spectrometry Facility, Texas A&M University, College Station, Texas, USA

**Keywords:** gram negative envelope, enterobacterial common antigen, O antigen, Wzz, glycan biosynthesis, chain length regulation

## Abstract

Enterobacterial common antigen (ECA) is a conserved glycan that supports outer membrane impermeability and intrinsic antibiotic resistance in Enterobacterales. ECA exists in outer membrane diacylglycerol-phosphate- and lipopolysaccharide-linked forms, and a cyclic periplasmic form (ECA_CYC_). Intriguingly, ECA_CYC_ both affects the outer-membrane permeability barrier and functions in regulation of diacylglycerol-phosphate-linked ECA levels. While the length of linear ECA polymers generated by WzyE is regulated by the co-polymerase WzzE, WzzE is also required for ECA_CYC_ biogenesis and no ECA_CYC_ is synthesized in its absence. To uncover WzzE functions necessary for ECA_CYC_ biosynthesis, we generated plasmid-borne and chromosomal *wzzE* mutants in *Escherichia coli* K-12 and quantified their effects on linear chain-length regulation and on ECA_CYC_ synthesis. Mutations affecting linear ECA chain-length regulation in either transmembrane helix 2 or the periplasmic domain abolished ECA_CYC_ synthesis. Notably, two variants at residue F104 exhibited identical linear ECA chain-length regulation which was similar to wild type, but resulted in sharply different ECA_CYC_ production: chromosomal WzzE_F104Y_ produced near wild-type levels of ECA_CYC_, whereas WzzE_F104H_ produced approximately two-fold less ECA_CYC_. This difference in ECA_CYC_ production vs. linear chain-length regulation demonstrates ECA_CYC_ synthesis is genetically separable from this regulation. Thus, WzzE-mediated chain-length regulation is necessary for ECA_CYC_ biogenesis, but not sufficient.

## Introduction

*Enterobacterales* is an order of gram-negative bacteria that includes numerous genera with human pathogens (e.g., *Escherichia, Salmonella, Klebsiella, Yersinia*) (Doern, 2024, Adeolu et al., 2016). These organisms cause a variety of diseases including respiratory, soft tissue, and urinary tract infections, as well as endocarditis (Ramirez and Giron, 2024). To survive in their enteric environment, members of this family take advantage of their outer membrane (OM) which acts as a robust permeability barrier composed of an asymmetric bilayer with phospholipids in the inner leaflet and predominantly lipopolysaccharides (LPS) in the outer leaflet (Maher and Hassan, 2023, Silhavy et al., 2010). LPS molecules form an extensive hydrophilic network of lateral interactions that excludes many large and hydrophobic compounds—such as many antibiotics, bile salts, and detergents—from entering the cell (Farfour et al., 2022, Aminov, 2010).

Beyond LPS, the outer membrane is decorated with other components critical for fitness, including enterobacterial common antigen (ECA) (Bennett and Mitchell, 2025, Rai and Mitchell, 2020). Identified in 1962 by Calvin M. Kunin et al. as a “common antigen” associated with urinary tract infections, ECA is an invariant carbohydrate restricted to *Enterobacterales* and absent only in endosymbionts with significantly reduced genomes (Rai and Mitchell, 2020, Jenkins et al., 2017, Kunin, 1963, Kunin et al., 1962). ECA has been shown to play a role in the pathogenesis in several members within the order of *Enterobacterales* (Aguiniga et al., 2016, Kalynych et al., 2014, Gilbreath et al., 2012, Armbruster et al., 2017, Holmes et al., 2023, Guérin et al., 2020). For example, *Salmonella enterica* serovar Typhimurium LT2 strains lacking ECA display reduced virulence, heightened sensitivity to bile salts, and establish a low-level persistent infection in mouse models (Kalynych et al., 2014, Gilbreath et al., 2012).

ECA is defined by recurring trisaccharide repeat units (RUs) of GlcNAc (*N*-acetyl-D-glucosamine), ManNAcA (*N*-acetyl-D-mannosaminuronic acid), and Fuc4NAc (4-acetamido-4,6-dideoxy-D-galactose) (Mannel and Mayer, 1978). There are three forms of ECA, (i) ECA_LPS_, which replaces O-antigen (Oag) on the LPS core; (ii) ECA_PG_, a cell-surface form linked to diacylglycerol-phosphate; and (iii) ECA_CYC_, a soluble, cyclic form found in the periplasm (Goździewicz et al., 2015, Kunin, 1963, Mannel and Mayer, 1978, Kajimura et al., 2005). ECA_PG_ and ECA_LPS_ vary in the number of RUs with a range of 1 to 14 and has a modal value of 5 to 7 at 37°C (Mitchell et al., 2018, Barr et al., 1999), whereas ECA_CYC_ has a defined number of exactly 4 RUs in *Escherichia coli* K-12 (Kajimura et al., 2005). Structural analysis studies including crystallography, nuclear magnetic resonance (NMR), and molecular dynamics indicate that ECA_CYC_ adopts either a rhomboidal or square conformations (Färnbäck et al., 2003, Erbel et al., 2004). We have previously found that ECA_CYC_ contributes to OM integrity in *E.coli* K-12 and mediates resistance to detergents and bile salts (Mitchell et al., 2018).

ECA biosynthesis genes reside mainly in the *wec* operon and employs a pathway analogous to the Wzy/Wzz-dependent pathway of Oag synthesis (Eade et al., 2021, Rick et al., 1985, Barr and Rick, 1987, Barr et al., 1989, Bertani and Ruiz, 2018, Barr et al., 1999, Erbel et al., 2003). After a single undecaprenyl-phosphate-linked ECA RU is synthesized, WzxE flips the intermediate to the outer leaflet of the inner membrane (Rai and Mitchell, 2020). WzyE and WzzE form a complex where the translocated undecaprenyl-phosphate-linked ECA subunits are polymerized by WzyE and the number of RU in the resulting chain is regulated by WzzE (Leo et al., 2021b, Maczuga et al., 2022b, Nath and Morona, 2015, Weckener et al., 2023). WzzE is an octameric inner-membrane protein with a large periplasmic domain (**Figure S1**) (Tocilj et al., 2008, Kalynych et al., 2012b, Collins et al., 2017, Kalynych et al., 2015). A cryo-EM structure from *Pectobacterium atrosepticum* suggest that WzyE resides within the central lumen formed by WzzE’s transmembrane helices (Weckener et al., 2023).

WzzE is a member of the class 1 polysaccharide co-polymerase (PCP1) family of chain length regulators, which includes WzzB, FepE, Wzz among others (Wiseman et al., 2021, Morona et al., 2009, Wiseman et al., 2023). Despite low sequence identity, PCP1 proteins share a conserved architecture consisting of two transmembrane helices and a bell-shaped periplasmic domain. (Tocilj et al., 2008, Kalynych et al., 2012b, Collins et al., 2017, Kalynych et al., 2015). In *Shigella flexneri*, WzzB C-terminal transmembrane helix two (TMH2) is known to interact with WzyB, the polymerase, and is necessary for Oag chain-length regulation (Ascari et al., 2022, Leo et al., 2021a). WzzB and WzyB co-purify, and the TMH2 of WzzE can maintain this interaction and contribute to Oag chain length regulation (Leo et al., 2021b). It has also been shown that mutations disrupting Oag chain-length regulation are found throughout Wzz and are not restricted to TMH2 (Whitfield et al., 2020).

WzzE plays a more complex role in ECA synthesis than other PCP1s do in Oag production. Loss of WzzE generates linear ECA with increased heterogeneity—producing ECA with both shorter and longer RUs—rather than the pronounced shortening typical of PCP1 deletions in Oag pathways (Leo et al., 2021b, Ascari et al., 2022, Huszczynski et al., 2019, Kalynych et al., 2011). In contrast to linear ECA, strains lacking WzzE fail to produce ECA_CYC_, indicating that WzzE is required for ECA_CYC_ synthesis despite a lack of known catalytic activity (Rai and Mitchell, 2020, Mitchell et al., 2018, Kajimura et al., 2005). In addition, WzzE plays a role in regulating levels of ECA_PG_ in *E. coli* K-12. ElyC regulates production of ECA_PG_ in such a way that loss of ElyC causes large increases in ECA_PG_ production (Rai et al., 2021). This regulation is dependent on WzzE and likely on ECA_CYC_, and changes in ECA_PG_ levels with loss of ElyC are much less in the absence of WzzE (Mitchell et al., 2018, Rai et al., 2021). Finally, ECA takes on aberrant activity in the absence of the anterograde phospholipid transporter, YhdP, causing increased sensitivity to vancomycin and the detergent SDS, and this phenotype is reversed when *wzzE* is absent (Mitchell et al., 2018).

Several non-exclusive hypotheses for how PCPs regulate chain length have been proposed (Islam and Lam, 2014, Leo et al., 2021b, Wiseman et al., 2021, Wiseman et al., 2023, Kintz and Goldberg, 2011, Papadopoulos and Morona, 2010, Morona et al., 1995, Weckener et al., 2023). (i) The inner or outer surface of the PCP may act as a molecular ruler for chain length regulation (Kintz and Goldberg, 2011). Specifically, the PCP may bind the extending carbohydrate, facilitating catalysis until the polymer length breaks the interaction (Kalynych et al., 2012a). (ii) The PCP could serve as a scaffold on which multiple polymerase molecules assemble (Tocilj et al., 2008). (iii) The elongating carbohydrate could extend in the lumen of the PCP until it reaches the top of the periplasmic domain and is exposed (Islam and Lam, 2014). (iv) The PCP could interact with the polymerase to sequester short glycan chains from downstream ligases (i.e., WaaL), increasing the processivity of polymerization (Hong et al., 2023). Finally, (v) the loops at the top of the periplasmic bell may move in an alternating up and down arrangement which allows for the polysaccharide chain to extend using a “ratchet”-type mechanism (Wiseman et al., 2023). However, the mechanism(s) used by WzzE for ECA_CYC_ synthesis is unknown. Here, we used site-directed mutagenesis and biochemical analysis to test whether WzzE requires only polymerase-interaction/chain-length regulation to produce ECA_CYC_, or if an additional function of WzzE contributes to ECA_CYC_ biogenesis. Our findings demonstrate that polymerase interaction and chain-length regulation are necessary for ECA_CYC_ biosynthesis. Moreover, we found that ECA_CYC_ synthesis genetically separated from linear ECA regulation, suggesting an additional function of WzzE in ECA_CYC_ synthesis.

## Results

### Plasmid-based *wzzE* allows analysis of *wzzE* mutant function

In order to define the functions of WzzE required for ECA_CYC_ synthesis in *E. coli* K-12, we examined the effects of *wzzE* mutations analogous to known residues affecting PCP1 Oag regulation. We first generated a set of *wzzE* constructs constitutively expressed by the *tet* promoter on the low copy number pZS21 plasmid (Lutz and Bujard, 1997), which does not contain a Tet repressor (**Figure 1A**). These included wild-type *wzzE* (p*wzzE*) and C-terminal fusions bearing a glycine-serine linker and triple FLAG tag (p*wzzE*-FLAG), a translation fusion to chloramphenicol acetyltransferase (p*wzzE-cat*), and a p*wzzE-cat* triple FLAG tag fusion (p*wzzE-cat-*FLAG). Based on the topology of WzzE, we anticipated that the C-terminal fusions would be exposed in the cytoplasm, thereby allowing Cat to function. Thus, these constructs allowed us to potentially identify defects in WzzE’s folding or stability as measured by chloramphenicol resistance. This approach was informed by a translational coupling methodology previously used a tool in *E. coli* (Mendez-Perez et al., 2012, Sieber, 2003), in which an unstable or degraded protein construct was shown to not allow chloramphenicol resistance. Additionally, we hypothesized that the formation of the functional cat trimer (Andreeva et al., 2000) is more likely if WzzE is properly multimerized, allowing Cat monomers to come into contact (with up to two Cat trimers potentially formed per octamer), giving insight into potential disruptions in octamerization. This assay is particularly useful because the addition of the bulky Cat (26KDa) protein likely imposes stress on WzzE folding. However, the chloramphenicol resistance results remain suggestive due to the potential limitations of this assay, such as the multimerization of Cat affecting octamer formation (i.e., a loss of chloramphenicol resistance is more indicative of a loss of stability and/or multimerization than chloramphenicol resistance is indicative of multimerization).

**Figure 1:**
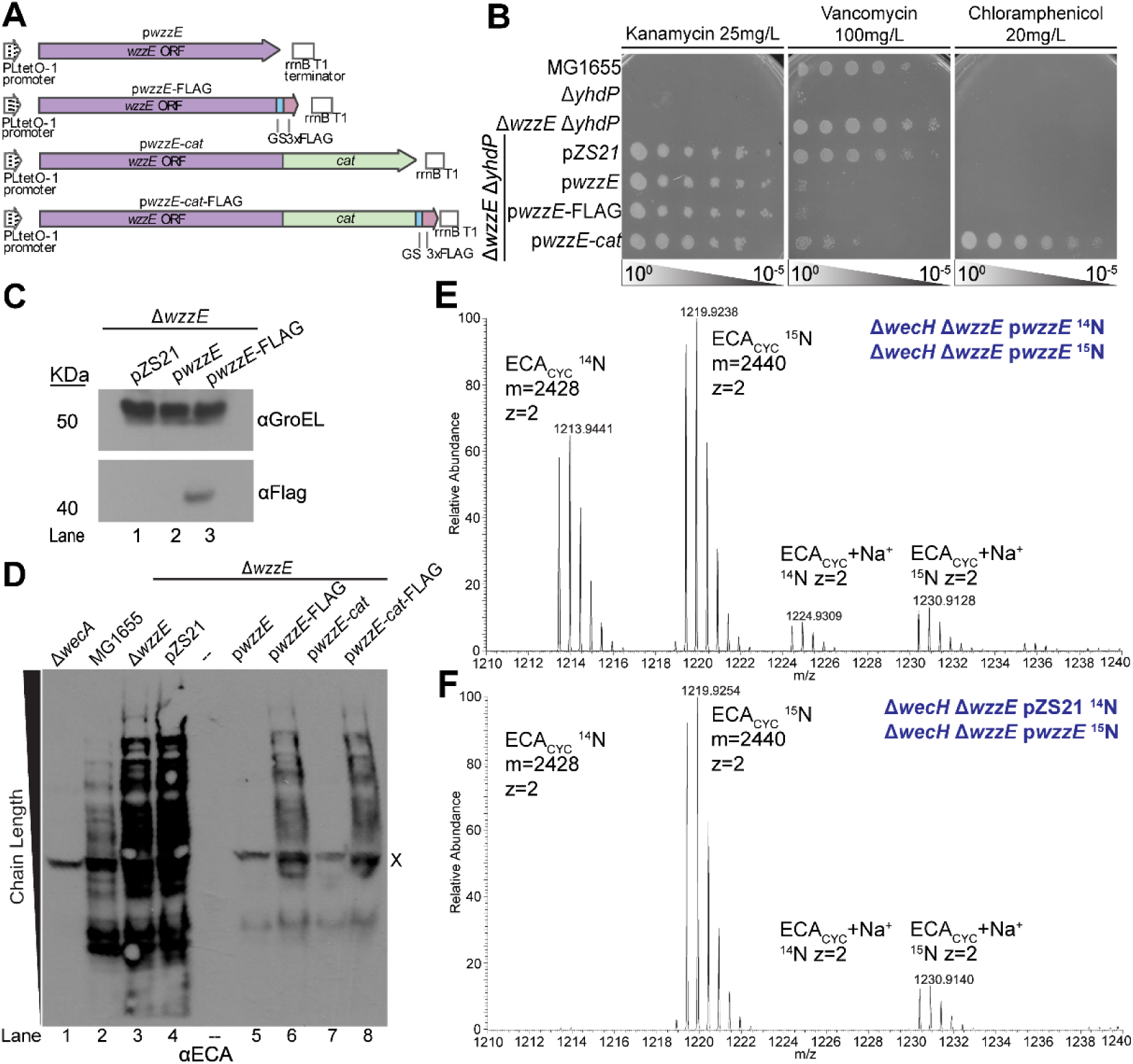
Plasmid-based expression of *wzzE* partially complements a chromosomal *wzzE* deletion. **(A)** Diagrams of constructs for expression of wild-type WzzE, WzzE with a C-terminal glycine-serine linker and triple FLAG tag, WzzE translationally fused with chloramphenicol acetyltransferase (Cat), and WzzE fused with Cat and triple FLAG are shown. The indicated constructs were expressed from the low copy number pZS21 plasmid (Lutz and Bujard, 1997). **(B)** The vancomycin sensitivity of a Δ*yhdP* strain can be suppressed by loss of WzzE (Mitchell et al., 2018). When the Δ*wzzE* Δ*yhdP* strain carried the pZS21 empty plasmid, it remained resistance to vancomycin as assayed by EOP but became sensitive when *wzzE, wzzE*-FLAG, or *wzzE*-*cat* was expressed, indicating all three constructs complemented Δ*wzzE*. In addition, translational fusion of *cat* and *wzzE* conferred chloramphenicol resistance, suggesting this construct could readout WzzE stability and multimerization based on chloramphenicol resistance. **(C)** WzzE-FLAG was detectable by immunoblotting. GroEL serves as a loading control. **(D)** ECA chain-length regulation was assayed through immunoblotting using an ECA antibody. Plasmid-base expression of WzzE restored chain-length regulation of linear ECA, although lower levels of ECA and shorter chain-length were observed than in the wild-type strain for the p*wzzE* and p*wzzE* constructs, while the p*wzzE-*FLAG and p*wzzE-cat-*FLAG constructs complemented better. The Δ*wecA* strain produces no ECA and serves as negative control. X: nonspecific band. **(E-F)** The strain carrying a plasmid with wild-type *wzzE* was grown with ^15^N to label ECA and then combined with an unlabeled strain carrying either wild-type *wzzE* (E) or the empty vector (F). ECA_CYC_ was purified and then detected by LC-MS. MS traces are shown for the indicated samples. The mass, charge, and identity of the ECA_CYC_-associated peaks are shown. Expression of *wzzE* from p*wzzE* restored ECA_CYC_ synthesis in the Δ*wzzE* strain. The strains contained Δ*wecH* to prevent non-stochiometric acetylation of ECA. All images are representative of at least three independent experiments.

We next assessed whether these constructs complemented Δ*wzzE.* In the absence of *yhdP*, ECA causes sensitivity to vancomycin in a *wzzE* dependent manner (Mitchell et al., 2018). Therefore, we expressed the *wzzE* constructs in a Δ*wzzE* Δ*yhdP* strain and tested for vancomycin sensitivity. Our results demonstrated each construct complemented Δ*wzzE* by restoring vancomycin sensitivity (**Figure 1B**). Additionally, p*wzzE*-*cat* conferred chloramphenicol resistance to the strain indicating that the Cat fusion was functional. To determine if the p*wzzE*-FLAG construct could be detected, we immunoblotted for FLAG using GroEL as a loading control (**Figure 1C**) and determined that the fusion protein could be clearly detected.

To determine whether linear ECA regulation was restored in the *wzzE* plasmid constructs, we first performed immunoblots for linear ECA (**Figure 1D**) using a polyclonal ECA antibody (Leo et al., 2021b). Our results revealed a range of ECA bands consistent with previous publications (Mitchell et al., 2018, Rai et al., 2021, Morris and Mitchell, 2023), with increasing molecular weight indicating molecules with more RUs. The distribution of ECA chain lengths was wider in a Δ*wzzE* mutant (Lane 3) than in wild-type cells (Lane 2) with an increase in longer and shorter ECA bands, indicating loss of chain-length regulation. We also observed more linear ECA with Δ*wzzE* as we would expect from previous findings (Rai et al., 2021, Mitchell et al., 2018). For simplicity, we will refer to these changes as “loss of chain-length regulation”. Expression of p*wzzE* (Lane 5) but not the empty vector (Lane 4) restored chain-length regulation, albeit with a lower modal chain length than the wild-type strain and much less linear ECA overall. These changes may suggest overactivity of WzzE due to multi-copy expression. The same pattern was observed for the p*wzzE-cat* construct (Lane 7); however, the p*wzzE*-FLAG and p*wzzE-cat-*FLAG constructs resulted in chain-length regulation closer to that of wild type.

Finally, to test whether the plasmid-based *wzzE* constructs rescue ECA_CYC_ synthesis, we expressed the p*wzzE* and empty vector plasmids in a Δ*wzzE* Δ*wecH* strain. WecH has been shown to non-stoichiometrically acetylate ECA (Kajimura et al., 2006) and deleting *wecH* allows ECA_CYC_ to be quantitated as a single mass species (Kajimura et al., 2006, Mitchell et al., 2018). Unlike the two linear forms of ECA, ECA_CYC_ is uncharged and cannot be detected by immunoblotting. Therefore, we grew cultures with either a normal (^14^N) or heavy (^15^N) nitrogen source, combined labeled and unlabeled samples, purified ECA_CYC_, and assayed ECA_CYC_ levels by liquid chromatography-mass spectrometry (LC-MS). ECA_CYC_ has a defined size of 4 RU in *E. coli* K-12 (Kajimura et al., 2005) and this molecule has an monoisotopic mass of 2428.9 Da in the absence of acetylation. Labeling with heavy nitrogen causes a 12 Da increase in this weight due to the presence of 12 nitrogen atoms in ECA_CYC_ (one in each sugar monomer) (Mitchell et al., 2018) (**Figure S2AB**). For this type of assay, each peak consists of multiple minor peaks due to the stochiometric incorporation of naturally occurring isotopes (e.g., ^14^C) (see **Figure 1EF**, **S2C**). When labeled and unlabeled samples carrying p*wzzE* were combined, we detected peaks at both the unlabeled and labeled m/z (**Figure 1E**). However, when the labeled p*wzzE* sample was combined with an unlabeled sample carrying the empty vector, only the ^15^N labeled ECA_CYC_ peak was detected (**Figure 1F**). ECA_CYC_ production from the p*wzzE* and p*wzzE-cat* constructs was very similar (**Figure S2C**). These data confirm plasmid-based expression of *wzzE* can facilitate the synthesis of ECA_CYC_, making this an appropriate system to investigate the role of WzzE in ECA_CYC_ biosynthesis.

### Conserved glycines in TMH2 are necessary for WzzE function

After validating our WzzE functional assays, we next examined the contribution of TMH2 on WzzE function. TMH2 has been shown to mediate interactions between WzzB and Wzy in O-antigen synthesis (Leo et al., 2021b), and prior mutagenesis studies of WzzB identified a conserved GXXXG motif in the TMH2 region of WzzB from *Shigella flexneri Y* that was important for function (Daniels and Morona, 1999), likely due to effects on helix-helix packing (Teese and Langosch, 2015, Prakash et al., 2011, Mueller et al., 2014). A corresponding double mutant in *Shigella flexneri Y* (G305A/G311A WzzB; G306A/G312A in *E. coli*) disrupts Oag chain length regulation (Papadopoulos et al., 2016); therefore, we hypothesized that the analogous mutations to GXXXG motif located in WzzE would play a significant role in mediating the helix-helix packing of the transmembrane region and its interaction with WzyE.

To test this, we generated TMH2 mutations (**Figure S1B**) in the p*wzzE* constructs described in **Figure 1A**. We first sought to determine whether the levels of WzzE protein produced by p*wzzE-*FLAG or p*wzzE-cat-*FLAG were altered by the mutations (**Figure 2AB, Figure S3A**). Our results showed a small but significant decrease in protein abundance for our WzzE^GG333/4LL^ in the WzzE-FLAG fusion (**Figure 2AB**), whereas no decrease is apparent in the WzzE-Cat-FLAG fusion (**Figure S3A**). As the levels of the WzzE^GG333/4LL^ variant are 82% of wild type on average and as high as 98% in individual replicates, the biological relevance of this decrease is unclear. To determine whether the mutations might have caused alterations to protein stability or multimerization, we utilized our p*wzzE-cat* construct to assay chloramphenicol resistance conferred by the fusion protein. None of our mutations showed any chloramphenicol sensitivity (**Figure 2C, Figure S3B**), suggesting that they are largely stable and multimerized, despite the small decrease in abundance for WzzE^GG333/4LL^. Next, we set out to determine if the mutations affected WzzE’s ability to regulate ECA chain length by performing ECA immunoblots. Three of the TMH2 mutations, WzzE^A323G^, WzzE^GG333/4AA^, and WzzE^G339A^ (Lanes 4,5, and 7) retained wild-type ECA chain length regulation (**Figure 2DE**). In contrast, WzzE^GG333/4LL^ (Lane 6) markedly disrupted chain-length regulation, resembling or exceeding the defect observed in our empty-vector control (Lane 3). These results identify the TMH2 GXXXG region as a key determinant of WzzE-mediated ECA chain-length regulation

**Figure 2:**
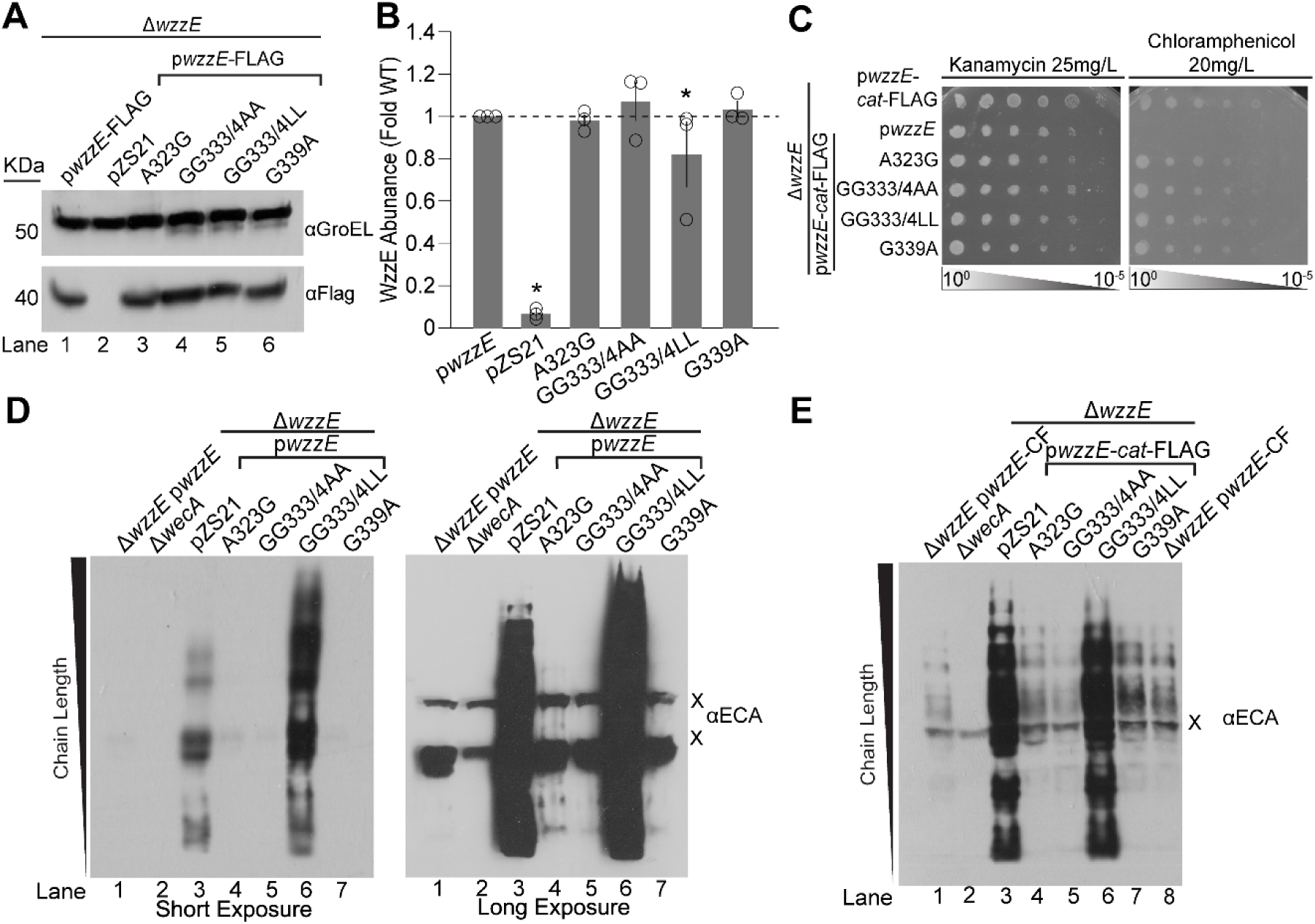
Alteration of WzzE TMH2 can prevent ECA chain length regulation without affecting WzzE stability. **(A)** The effect of the indicated *wzzE* site-directed mutations on WzzE-FLAG protein levels was assessed using immunoblotting. GroEL serves as a loading control. **(B)** Densitometry was used for quantification of the immunoblots in (A). Levels of WzzE-FLAG compared to GroEL are shown normalized to the wild-type *wzzE* complemented strain. All variants show similar protein levels to wild-type WzzE, except for a small, but significant decrease in the WzzE^GG333/4LL^ variant. Data are shown as the average of three biological replicates ± SEM and individual datapoints. * p<0.05 by Mann-Whitney test. **(C)** Strains carrying the indicated plasmids were assayed for their chloramphenicol resistance by EOP as a proxy for WzzE stability and multimerization ability. All *wzzE* mutants retained full chloramphenicol resistance, suggesting WzzE stability and multimerization. **(D)** Chain-length regulation of the linear ECA forms by the indicated WzzE variants was assayed by immunoblotting. Short and long exposures are shown. “X” indicates non-specific bands. WzzE^GG333/4LL^ showed loss of chain-length regulation while the other variants retained wild-type function. The Δ*wecA* strain serves as negative control. **(E)** Chain-length regulation of the WzzE-Cat-FLAG variants was assayed by immunoblot. Similar phenotypes were observed to those found in the WzzE variants. “X” indicates non-specific band. All images are representative of at least three independent experiments.

### The TMH2 GXXXG motif affects ECA_CYC_ synthesis

Given the strong defect of WzzE^GG333/4LL^ in regulating ECA chain length, we assessed whether the TMH2 mutations impair ECA_CYC_ biogenesis. We utilized LC-MS to detect and quantitate ECA_CYC_ in purified samples as in **Figure 1EF**. We expressed the TMH2 mutants or controls in a Δ*wzzE* Δ*wecH* strain background and labeled the wild type p*wzzE* strain with ^15^N, while the mutants were grown with ^14^N. The quantification demonstrated WzzE^A323G^, WzzE^GG333/4AA^, and WzzE^G339A^ did not significantly change ECA_CYC_ levels, whereas the WzzE^GG333/4LL^ mutant produced significantly reduced levels of ECA_CYC_, similar to those of the empty vector control (**Figure 3A**). These data demonstrate that the TMH2-dependent chain-length regulation is required for ECA_CYC_ synthesis.

**Figure 3.**
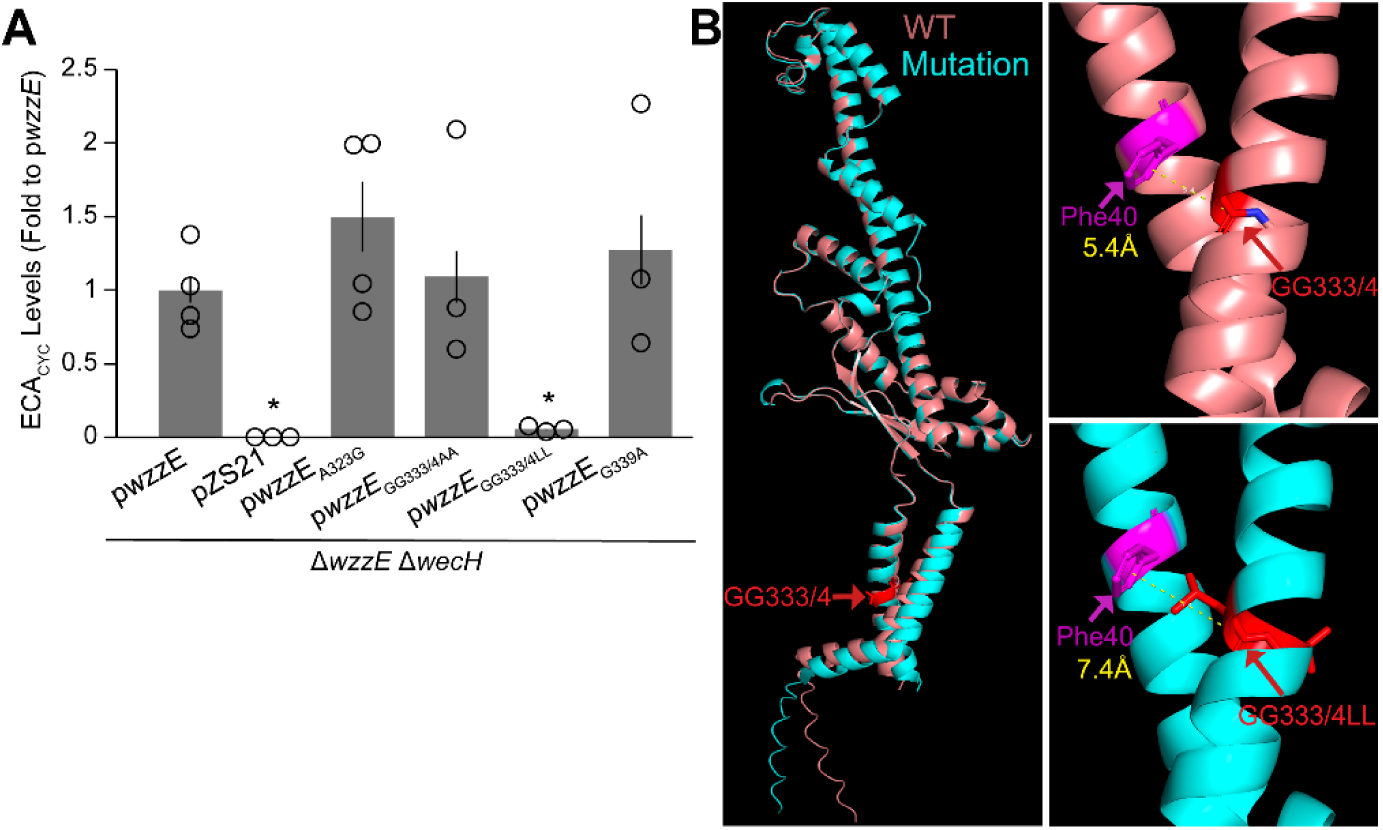
Conserved glycines in WzzE TMH 2 are required for ECA_CYC_ synthesis. **(A)** The strain carrying a plasmid with wild-type *wzzE* was grown with ^15^N to label ECA and then combined with an unlabeled strain carrying the indicated *wzzE* mutant or control. ECA_CYC_ was purified and the relative amount of unlabeled to labeled ECA_CYC_ was determined by LC-MS. WzzE^GG333/4LL^ showed a loss of the ability to produce ECA_CYC_. All plasmids were expressed in a Δ*wzzE* Δ*wecH* strain background to prevent non-stochiometric acetylation of ECA. Data are shown as the average of three biological replicates ± SEM and as individual datapoints. * p<0.05 by Mann-Whitney test. **(B)** The predicted structure of a WzzE octamer with WzyE was calculated using AlphaFold 3 (Abramson et al., 2024). The structures of wild-type WzzE and WzzE^GG333/4LL^ monomers were aligned with the mutated residues shown in red. The insets show the predicted distance between the two WzzE TMH in the wild type WzzE and WzzE^GG333/4LL^.

To visualize how these mutations might alter WzzE function, we modeled the WzyE:WzzE(8) complex for wild-type WzzE and WzzE^GG333/4LL^ using AlphaFold 3 (Abramson et al., 2024) (**Figure 3B** & **S4**, **Table S1**). Alignment of monomers revealed a ∼2 Å expansion between TMH1 and TMH2 in WzzE^GG333/4LL^ consistent with steric disruption of helix-helix packing (**Figure 3B**). This structural shift provides a potential mechanistic basis for the loss of both chain-length regulation and ECA_CYC_ biosynthesis in our GXXXG mutant.

### The periplasmic domain of WzzE functions in chain length regulation

After assessing the importance of WzzE TMH2 for ECA_CYC_ biosynthesis, we next sought to determine whether chain-length regulating activities within the periplasmic domain of WzzE are also required for ECA_CYC_ biosynthesis. Twenty-one periplasmic residues are conserved among WzzE, FepE, and WzzB in *E. coli*, suggesting that they are important for PCP1 function (Tocilj et al., 2008). We also hypothesized that non-conserved residues could contribute to WzzE-mediated chain-length regulation or ECA_CYC_ synthesis. Therefore, we made mutations in a conserved cluster of predominantly hydrophobic residues located in the core of the ⍺/β domains—spanning strands β1 and β4, helix ⍺2 and the N terminus of helix ⍺6 (see **Figure S1B**). Additionally, we targeted non-conserved residues throughout WzzE in positions similar to those that conferred changes to Oag chain length in other PCP1s (Tocilj et al., 2008).

We first characterized the periplasmic mutants (**Figure S1B**) by examining WzzE protein levels using the p*wzzE*-FLAG construct. WzzE^ΔF104^ (an in-frame, single codon deletion) had undetectable WzzE-FLAG protein levels, whereas all other mutants retained wild-type protein accumulation (**Figure 4AB**). When we examined the levels of WzzE-Cat-FLAG for WzzE^ΔF104^, we observed a faint band at the expected molecular weight of the fusion and a second, lower-molecular-weight band, indicating cleavage of the C-terminal tag (**Figure S3C**). Given the cleavage of the tags, no conclusion could be drawn for the stability of WzzE^ΔF104^. All other protein levels were similar to those of wild type.

**Figure 4:**
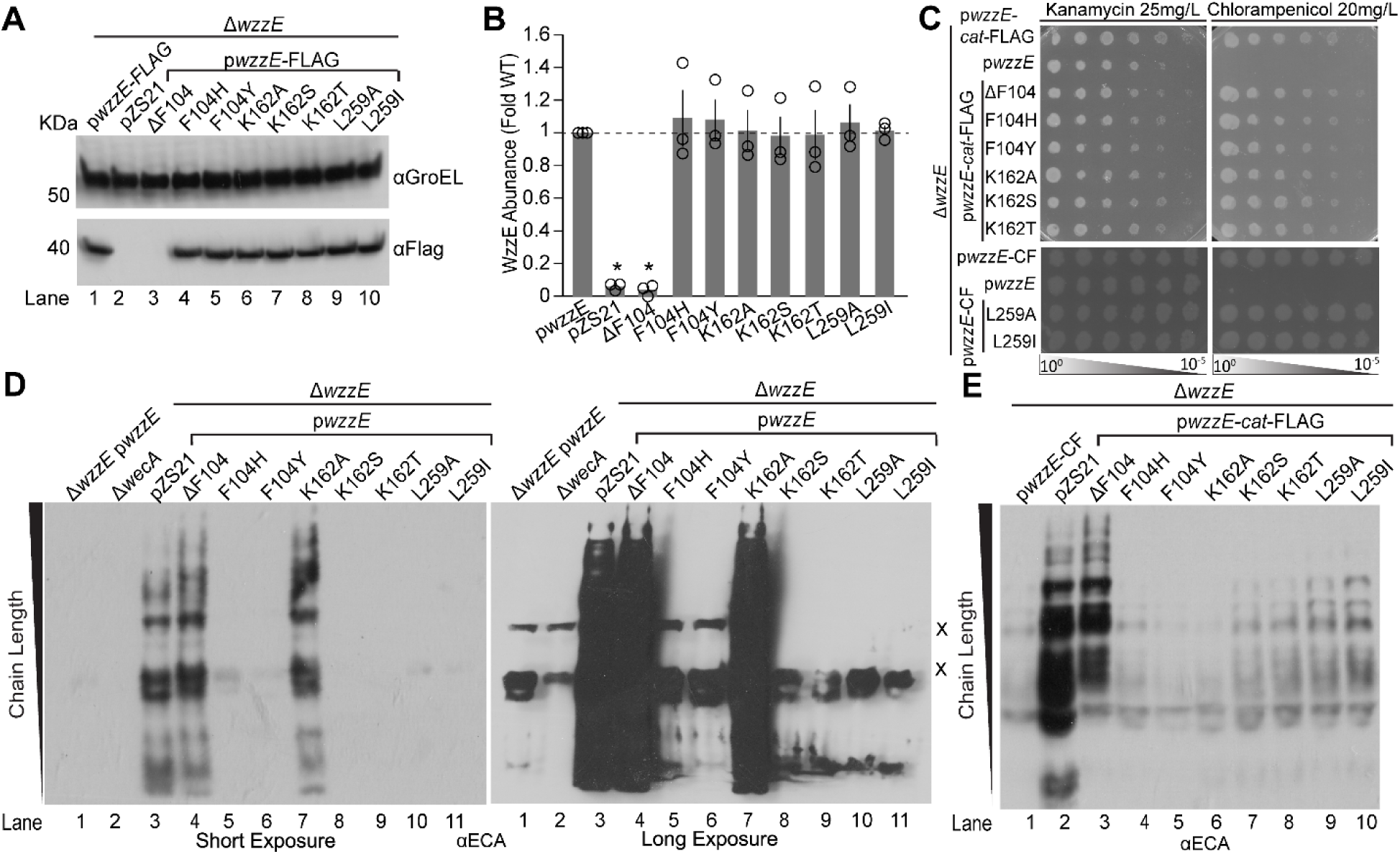
The periplasmic region of WzzE contributes to WzzE stability and chain length regulation. **(A)** Immunoblotting was used to determine protein levels for the indicated WzzE-FLAG variants. GroEL serves as a loading control. **(B)** WzzE-FLAG levels from (A) were quantified by densitometry. WzzE-FLAG levels compared to GroEL are shown normalized to the wild-type *wzzE* complemented strain. All variants are similar to wild type WzzE except for WzzE^ΔF104^ which shows significantly reduced protein levels. Data are shown as the average of three biological replicates ± SEM and individual datapoints. * p<0.05 by Mann-Whitney test. **(C)** The indicated strains were assayed for WzzE-Cat-FLAG stability as measured by chloramphenicol resistance. All *wzzE* point mutations showed chloramphenicol resistance indicating stability and multimerization, with the exception of WzzE^ΔF104^ where the tag is cleaved (**Figure S3C**). **(D)** Linear ECA chain-length regulation by WzzE variants was assayed by immunoblotting with ECA antibody. Long and short exposures are shown. The periplasmic point mutants have variable effects on chain length regulation as well as ECA_CYC_ biogenesis. The WzzE^ΔF104^ and WzzE^K162A^ variants showed loss of ECA chain-length regulation while the other variants had wild-type chain-length regulation. The Δ*wecA* strain serves as negative control. **(E)** The effect of WzzE-Cat-FLAG variants on ECA chain-length regulation was assayed by immunoblot. The variants show similar phenotypes as in (D) except for WzzE^K162A^, which shows normal chain-length regulation and WzzE^ΔF104^ which shows partial chain-length dysregulation. All images are representative of at least three independent experiments.

We next evaluated whether the *wzzE* mutations might affect protein stability and/or multimerization by measuring chloramphenicol resistance conferred by the p*wzzE*-cat fusion. WzzE^K162A^, WzzE^L259A^, and WzzE^F104Y^ caused partial or full chloramphenicol sensitivity in the p*wzzE*-Cat fusion (**Figure S3D**) but showed full chloramphenicol resistance in the p*wzzE*-Cat-FLAG fusion, which better complements linear chain length regulation (**Figure 4C**). These differences suggest these variants may exhibit mild instability or reduced multimerization. All other mutants conferred chloramphenicol resistance similar to wild type in both fusions (**Figure 4C, S3D**). WzzE^ΔF104^ showed wild-type levels of chloramphenicol resistance, likely reflecting cleavage of the C-terminal tag. We also tested an additional 18 periplasmic-domain mutations that caused chloramphenicol sensitivity in the WzzE-Cat background, emphasizing the importance of this domain for WzzE structural stability and multimerization (**Figure S5**).

After characterizing the effect of the periplasmic mutations on WzzE levels and stability, we next examined how these periplasmic mutations affect ECA chain-length regulation (**Figure 4DE**). WzzE^ΔF104^ and WzzE^K162A^ (lanes 4 and 7) disrupted chain-length regulation when expressed from p*wzzE* (**Figure 4D**). However, when expressed from p*wzzE*-cat-Flag, WzzE^ΔF104^, exhibited partial impairment of chain-length regulation, while WzzE^K162A^ showed wild-type chain-length regulation (**Figure 4E**). These differences parallel the chloramphenicol resistance phenotypes observed for the corresponding fusions (**Figure 4C**, **S3D**). In contrast, WzzE^F104Y^, WzzE^F104H^, WzzE^K162S^, WzzE^K162T^, WzzE^L259A^, and WzzE^L259I^ (lanes 5, 6, 8-11) demonstrated wild-type chain length regulation in both the p*wzzE* and p*wzzE-cat-FLAG* constructs. Together, these mutants provide a useful range of phenotypes for dissecting the requirements for ECA_CYC_ synthesis.

### Periplasmic residues of WzzE are important for ECA_CYC_ biogenesis

We next assessed the impact of our periplasmic mutants on ECA_CYC_ biosynthesis by quantifying ECA_CYC_ levels using LC-MS. Both the WzzE^ΔF104^ and WzzE^K162A^ mutants exhibited impaired linear chain-length regulation and produced little to no ECA_CYC_ (**Figure 5A**). This loss of ECA_CYC_ production could results from either (i) insufficient protein accumulation and/or stability or (ii) disruption of chain-length-regulating activities in these mutants. In contrast, WzzE^F104Y^, WzzE^K162S^, WzzE^K162T^, WzzE^L259A^, and WzzE^L259I^ produced ECA_CYC_ at levels comparable to wild type. WzzE^F104H^ showed highly variable ECA_CYC_ levels despite maintaining wild type chain-length regulation and protein levels, suggesting that this mutation may have a specific ECA_CYC_ synthesis effect. These data highlight the importance of WzzE’s periplasmic domain on ECA_CYC_ synthesis, likely through its structural role, its chain-length-regulating activity, and possibly a function specific to ECA_CYC_ synthesis. Given the differing effects of the F104 mutations on ECA_CYC_ synthesis and linear ECA chain-length regulation, we examined their potential structural consequences using AlphaFold 3 (Abramson et al., 2024) and aligned wild type and WzzE^ΔF104^ complexes (**Figure 5B** & **S4**, **Table S1**). These models predict that F104 forms contacts with a neighboring loop, and alterations in these interactions may underlie the functional differences observed for substitutions at this residue.

**Figure 5:**
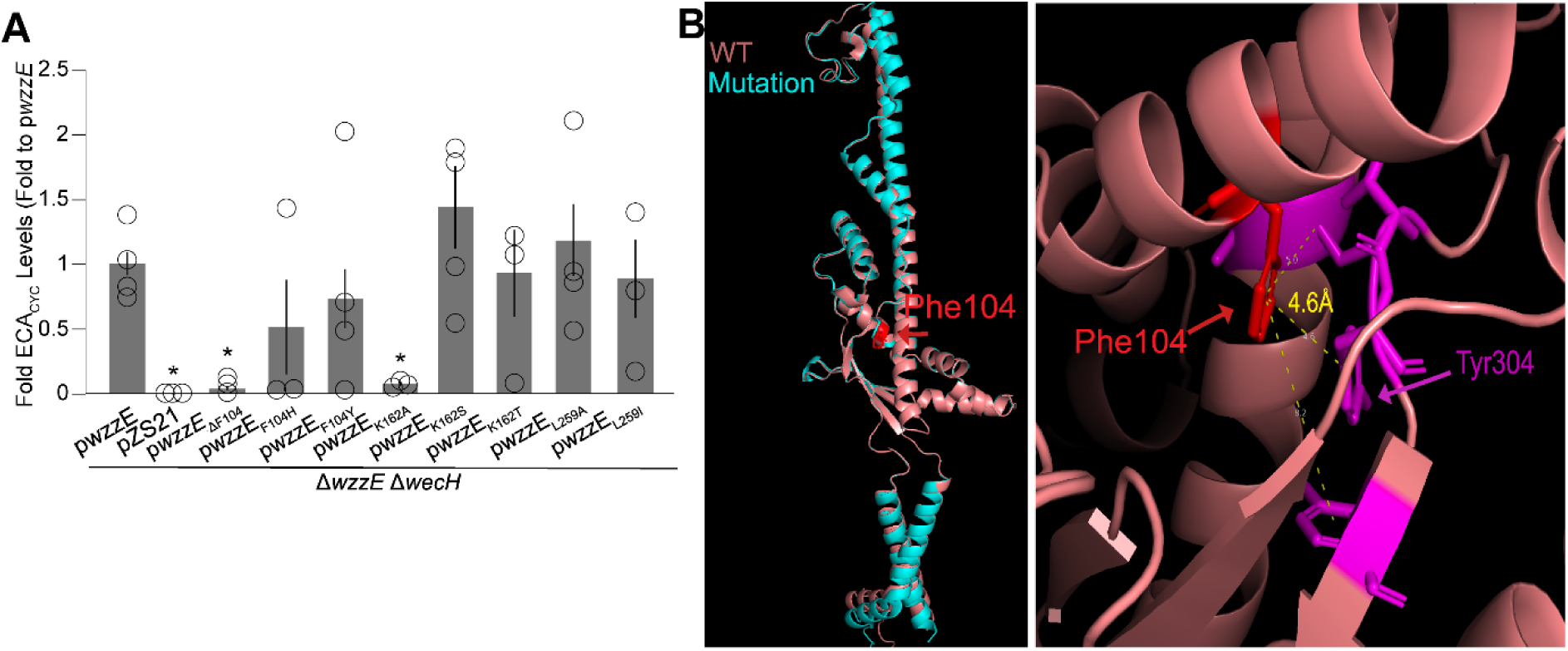
The periplasmic domain of WzzE contributes to ECA_CYC_ synthesis. **(A)** The strain carrying wild-type *wzzE* was grown with ^15^N to label ECA and then combined with an unlabeled strain carrying the indicated *wzzE* mutant or control strain. After ECA_CYC_ purification, the relative amount of unlabeled to labeled ECA_CYC_ was determined by LC-MS. WzzE^ΔF104^ and WzzE^K162A^ had little to no ECA_CYC_ production while WzzE^F104H^ showed a variable phenotype. All strains included Δ*wecH* to prevent non-stochiometric acetylation of ECA. Data are shown as the average of three biological replicates ± SEM and as individual datapoints. * p<0.05 by Mann-Whitney test. **(B)** Structures predicted as in Figure 3 for wild-type WzzE and WzzE^ΔF104^ were aligned. The mutated residue is shown in red on a WzzE monomer (B). The inset shows the interaction between alpha helix 2 and an adjoining loop in wild-type WzzE.

### ECA_CYC_ biogenesis is genetically separable from chain-length regulation

Our plasmid-based expression analyses of *wzzE* identified several point mutations with distinct effects on ECA_CYC_ production. Notably, substitution of the TMH2 residues (WzzE^GG333/4LL^) abolished chain-length regulation and ECA_CYC_ synthesis, while maintaining protein stability and near-wild-type protein accumulation. Additionally, several mutations to residue F104 also produced variable effects on WzzE accumulation, chain-length regulation, and ECA_CYC_ production. However, plasmid-based expression of wild-type *wzzE* yielded lower levels of linear ECA and reduced modal chain-length compared expression of *wzzE* from its native locus (**Figure 1D**), a difference likely contributing to some of the phenotypes observed in the plasmid system. We also noted substantial variability in ECA_CYC_ levels among periplasmic mutants, which we attributed to non-native gene expression. To circumvent these limitations, we used CRISPR-Cas9 to introduce select point mutations into the native *wzzE* locus and assayed their effect on linear ECA and ECA_CYC_.

LC-MS analysis of purified ECA_CYC_ revealed that chromosomal WzzE^ΔF104^ and WzzE^GG333/4LL^ mutants produced little to no ECA_CYC_, similar to what is observed with Δ*wzzE* (**Figure 6A**). In contrast, WzzE^F104H^ produced measurable levels of ECA_CYC_ that were about 2-fold lower than the wild type, whereas WzzE^F104Y^ showed no significant change in ECA_CYC_ levels relative to the wild type. We then assayed linear ECA chain length regulation by immunoblot (**Figure 6B**). Consistent with the plasmid-based system, WzzE^ΔF104^ and WzzE^GG333/4LL^ failed to regulate linear ECA. The magnitude of chain-length dysregulation in these mutants exceeded that of Δ*wzzE*, potentially suggesting an aberrant interaction of non-functional WzzE and ECA synthesis that differs from the absence of WzzE. WzzE^F104H^ and WzzE^F104Y^ each displayed intact and nearly wild-type chain-length regulation (**Figure 6B**), with a possible slight increase in modal chain length that was not present in all replicates. However, the linear ECA regulation of WzzE^F104H^ and WzzE^F104Y^ was consistently identical between these two mutants.

**Figure 6:**
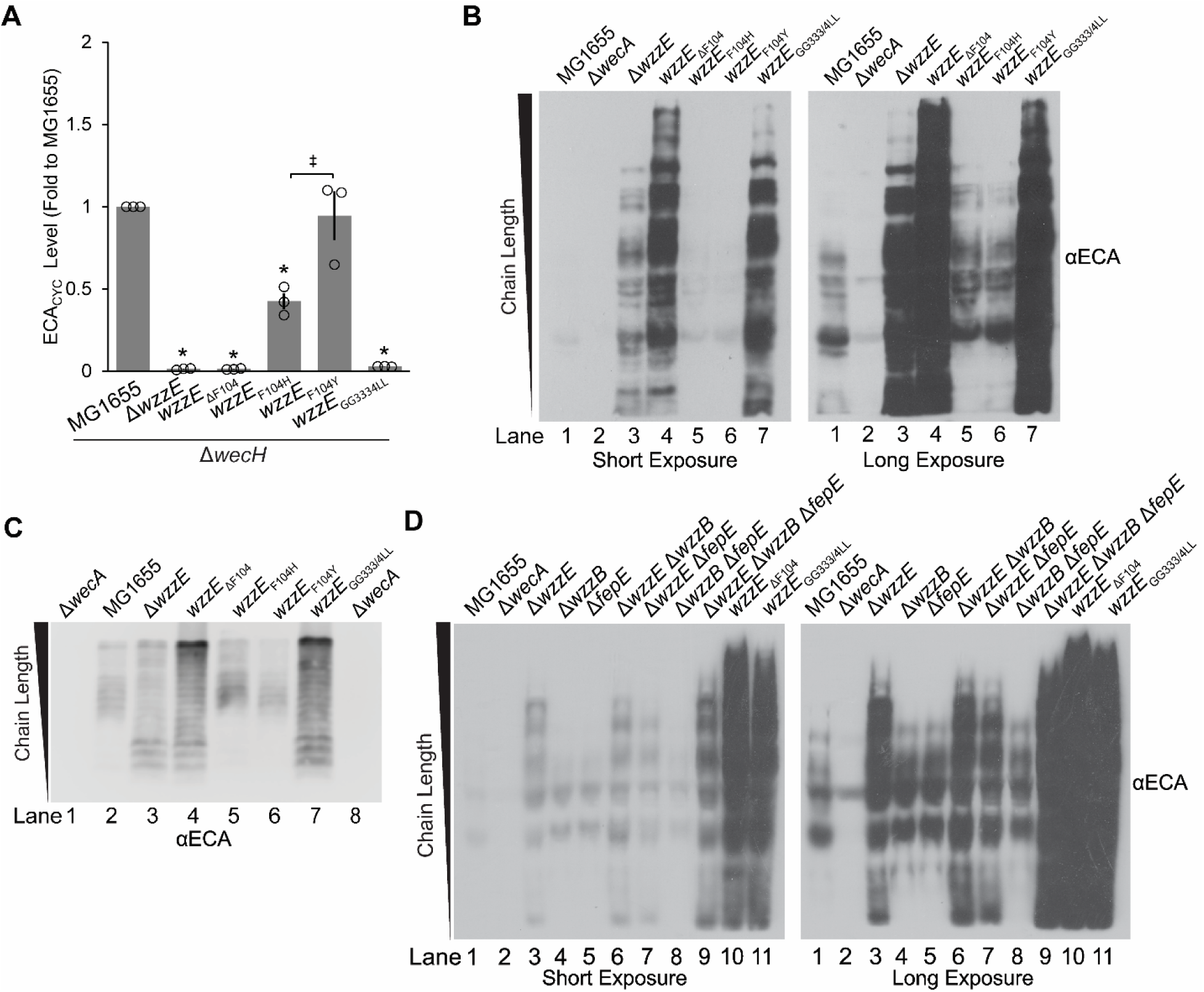
Levels of ECA_CYC_ production and chain-length regulation are genetically separable. CRISPR-Cas9 was used to make the indicated mutations to *wzzE* in its native locus. All strains contain *metE*::Tn*10*. **(A)** The strain wild type for *wzzE* was grown with ^15^N to label ECA and then combined with an unlabeled strain with the indicated *wzzE* mutant or control strain. After ECA_CYC_ purification, the ratio of unlabeled to labeled ECA_CYC_ was determined by LC-MS. All variants other than WzzE^F104Y^ produced significantly reduced ECA_CYC_ levels. Data are shown as the average of three biological replicates ± SEM and as individual datapoints. * p<0.05 vs. wild type by Mann-Whitney test; ‡ p<0.05 between the indicated strains by Mann-Whitney test. **(B)** Immunoblotting for ECA was conducted on the indicated chromosomal mutants. A short and long exposure are shown. The WzzE^ΔF104^ and WzzE^GG333/4LL^ variants resulted in linear ECA dysregulation, while the other mutations more closely resembled wild-type linear ECA regulation. The Δ*wecA* strain serves as negative control. **(C)**. ECA Immunoblotting was performed on the indicated chromosomal mutants after ProK treatment to more clearly assess chain length. WzzE mutants ΔF104 and GG333/4LL resulted in dysregulation of linear ECA, whereas F104Y and F104H strongly resembled wild-type linear ECA regulation. **(D)** Immunoblotting was used to assay ECA chain-length regulation in the indicated strains. Deletion of *wzzB* and/or *fepE* in combination with Δ*wzzE* did not cause the increased levels of linear ECA observed with the loss-of-function point mutants; however, the Δ*wzzE* Δ*wzzB* Δ*fepE* strain did have a small increase in linear ECA levels compared to Δ*wzzE* alone. All images are representative of at least three independent experiments.

In our ECA immunoblot protocol, we take steps to minimize ECA degradation (see Experimental Procedures), as ECA can be delipidated by mild acid hydrolysis (BASU et al., 1987). One of these steps is to avoid long incubations, such as those used to proteinase K (ProK) treat LPS samples (May and Silhavy, 2018). However, protein in our samples cause ECA to spread over a larger area of the gel and potentially could mask small changes in chain length regulation. To improve detection of small changes in chain length while minimizing ECA degradation, we pretreated our immunoblot samples with various concentrations of ProK and incubation times at 56 °C found in the literature (Leo et al., 2020, Jiang et al., 2020, Maczuga et al., 2022a, Liu et al., 2020, Peters et al., 1985), as well as treatments optimized based on our initial results. The resulting blots and comparative signal quality are shown in (**Figure S5**). We selected a one-hour treatment with a high concentration of ProK to move forward with as this treatment resulted in loss of protein signal while minimizing ECA degradation. After treatment with ProK in these optimized conditions, both WzzE^F104H^ and WzzE^F104Y^ showed ECA chain length that was extremely similar to wild type (**Figure 6C**). Overall, the identical regulation of linear ECA chain-length by WzzE^F104H^ and WzzE^F104Y^ but differing effects on ECA_CYC_ synthesis demonstrate that levels of ECA_CYC_ production are genetically separable from linear chain-length regulation.

We were surprised that the dysregulation of ECA chain length, and even more the levels of ECA, in the WzzE^ΔF104^ and WzzE^GG333/4LL^ mutants was still greater than the Δ*wzzE* control, and so we wondered whether, in a Δ*wzzE* strain, the remaining PCP1s could partially regulate ECA chain length through interaction with WzyE. In this model, nonfunctional WzzE mutants might sequester WzyE from the other PCP1s, exacerbating dysregulation. Therefore, we assayed whether we could cause further disruption of ECA chain-length regulation in a Δ*wzzE* strain by deleting *wzzB* and/or *fepE*. Deletion of Δ*wzzB*, Δ*fepE*, and Δ*wzzB* Δ*fepE*, or both had no detectable effect on chain regulation in a Δ*wzzE* strain (**Figure 6D**), though the triple mutant showed a modest increase in linear ECA levels. Taken together, these findings demonstrate that mutations in *wzzE* that disrupt linear ECA chain-length regulation can also abolish ECA_CYC_ synthesis, and that normal ECA_CYC_ production is genetically separable from regulation of linear ECA chain-length.

## Discussion

In this study, we constructed and characterized a set of *wzzE* site-directed mutants to define which functions of WzzE are required for ECA_CYC_ biogenesis. Although previous mutagenesis studies have identified regions of *Shigella flexneri* WzzB important for activity (Nath and Morona, 2015, Ascari et al., 2022), no prior work has examined how WzzE contributes to the production of cyclic ECA. Here, we show that substitutions at F104 within the periplasmic domain and to the GXXXG motif in TMH2 can compromise both ECA chain-length regulation and ECA_CYC_ synthesis. Moreover, our data demonstrate that regulation of linear ECA chain length and the level of ECA_CYC_ production are genetically separable, as we have identified two mutations at F104 with divergent effects on ECA_CYC_ levels, but produce identical regulation of linear ECA chain length. These data show regulation of chain length by WzzE is necessary but not sufficient for production of normal levels of ECA_CYC_.

Despite low sequence conservation among PCP1s, their overall structures are highly similar, consisting of two transmembrane domains and an ⍺-helix/β-sheet domain within the periplasmic space that extends ∼100 Å above the membrane (Tocilj et al., 2008, Kalynych et al., 2012b, Collins et al., 2017, Kalynych et al., 2015). PCP1s also harbor a conserved GXXXG motif that has been shown to contribute to chain-length regulation. In WzzB, a G305A/G311A double mutant in *Shigella flexneri* WzzB that encompassed and extended outside of this motif resulted in short-chain O-antigen (Oag) (Daniels and Morona, 1999, Papadopoulos et al., 2016), whereas single, double, and triple glycine to alanine mutations restricted to the GXXXG motif did not affect chain-length (Collins et al., 2017, Daniels and Morona, 1999). Motivated by these findings, we focused on residues within and around the GXXXG motif, characterized their effect on chain-length regulation and ECA_CYC_ biogenesis, and identified two adjacent, conserved glycine residues in the GXXXG motif of WzzE—analogous to G305/G306 in WzzB—as critical for ECA chain-length regulation and ECA_CYC_ biosynthesis. Interestingly Ala substitutions at these positions had no detectable effect on chain-length regulation or ECA_CYC_ production, whereas introduction of adjacent Leu residues led to pronounced dysregulation of ECA chain-length and significantly reduced levels of ECA_CYC_.

GXXXG motifs often mediate dimerization and multimerization of transmembrane domains (Parrish et al., 2015, Faingold et al., 2012), suggesting that the motif may facilitate positioning of TMH1 and TMH2 of WzzE. In support of this idea, our structural modeling suggests increased distance between TMH1 and TMH2 in the WzzE^GG333/4LL^ mutant (**Figure 3D**, **S6A**). Alignments of WzzE monomers predicted to closely interact with WzyE, shows that WzyE rotates in the plane perpendicular to the membrane in WzyE-WzzE^GG333/4LL^ model compared to the wild type (**Figure S6BC**). The alignment also shows the distance in the structure between the TMH of opposite WzzE monomers was 2.0 to 2.7 Å smaller in the WzzE^GG333/4LL^ model than in the wild-type model. In compensation, WzyE takes on a more condensed structure (1.4 Å narrower) in the WzzE^GG333/4LL^ model. As this type of rotation and structure change could impair WzyE function, this prediction provides a possible hypothesis that WzzE^GG3334/LL^ has impaired interaction with WzyE. Alternatively, disruption of TMH packing may interfere with an intrinsic and lesser defined activity of WzzE.

In contrast to mutations in TMH2, many substitutions introduced into the periplasmic domain rendered WzzE unstable, as evidenced by loss of chloramphenicol resistance conferred by the WzzE-Cat fusion. This observation underscores the importance of the periplasmic domain for WzzEs structural integrity and/or multimerization. Prior work on other PCP1s suggests that multiple surface-exposed residues within the periplasmic domain affect chain-length regulation, producing either shorter or longer Oag chains (Kalynych et al., 2012b, Papadopoulos and Morona, 2010, Daniels and Morona, 1999, Hong and Payne, 1997, Tran and Morona, 2013). More recently, increasing evidence suggests that the inner surface of the WzzE octamer may bind to the elongating carbohydrate to facilitate growth of the chain (Wiseman et al., 2023, Hong et al., 2023, Weckener et al., 2023).

WzzE’s internal cavity possesses two highly concentrated, negatively charged bands, separated by a flexible neutrally charged loop that may bind the growing carbohydrate chain (Wiseman et al., 2023). A loop at the apex of the periplasmic domain has been suggested to “ratchet” up and down to facilitate carbohydrate movement over this binding surface (Wiseman et al., 2023). F104 is buried near the base of the periplasmic “bell” and may affect this binding surface indirectly. In contrast, K162 is positioned adjacent to the lower negatively charged band, and L259 residues within the proposed ratchet loop. Mutations to L259 did not affect chain-length regulation; however, loss of charge at K162 did prevent chain-length regulation and ECA_CYC_ synthesis, perhaps supporting this model of ECA chain-length elongation. As to how the *wzzE_K162A_* and *wzzE _ΔF104_* mutations lead to loss of ECA_CYC_ production, we speculate that the chain-length regulating activity of WzzE—related to the carbohydrate binding and ratcheting mechanism—is necessary for ECA_CYC_ synthesis.

In contrast, the *wzzE_F104H_* mutation reduces ECA_CYC_ levels while maintaining comparatively wild-type chain-length regulation. Moreover, its chain-length regulation is indistinguishable from *wzzE_F104Y_*, which produces wild-type ECA_CYC_ levels. These results demonstrate that chain-length regulation alone is insufficient for normal ECA_CYC_ synthesis and suggest that WzzE plays an additional role specific to ECA_CYC_ synthesis beyond that of chain-length regulation. One possible mechanism would be that WzzE bends ECA molecules of the correct size into an orientation that can be spontaneously cyclized. Another possibility is that WzzE allows the binding of another protein (i.e., a cyclase) that can cyclize ECA chains of the appropriate length. Finally, it is possible that WzzE has specific features that promote the formation of ECA chains with 4 RUs that can later be cyclized to form ECA_CYC_. Further, genetic and biochemical experiments will be necessary to distinguish between these possibilities and to solidify an additional role for WzzE in ECA_CYC_ biogenesis.

Unexpectedly, we consistently observed that WzzE ^ΔF104^ and WzzE^GG333/4LL^ mutants had greater linear ECA levels than what was observed in our *wzzE* deletion. We confirmed that this phenotype does not arise from compensation by other PCP1s in *E.coli*, although we did notice a modest increase in linear ECA in strains lacking all other PCP1 genes. Therefore, why WzzE ^ΔF104^ and WzzE^GG333/4LL^ have elevated linear ECA levels remains unclear. One explanation is that inactive WzzE octamers partially sequester growing ECA chains, increasing variability in chain length. Regardless, we believe that the phenotypic distinction between Δ*wzzE* strain and the mutants strengthens the conclusion that these mutants produce inactive proteins which accumulate but are incapable of regulating chain-length or supporting ECA_CYC_ biosynthesis.

Overall, our analysis interrogates only a small fraction of WzzE residues, and additional mutations that selectively impair ECA_CYC_ biosynthesis likely exist. Identification of such mutants will clarify the molecular mechanism of ECA_CYC_ biosynthesis. Nonetheless, our findings reveal fundamental requirements for ECA_CYC_ and chain-length regulation. We propose coordinated activity between WzyE and WzzE, together with periplasmic charge elements within WzzE that facilitate chain elongation, is required for ECA_CYC_ biogenesis. Importantly, we demonstrate that chain-length regulation can be genetically uncoupled from ECA_CYC_ synthesis, indicating that WzzE possesses a second function beyond chain-length regulation that is important for ECA_CYC_ biosynthesis. These insights help to define key features of WzzE that enable synthesis of a cyclic carbohydrate.

## Experimental Procedures

### Strains and growth conditions

The strains used in this study are listed in **Table S2**. Cultures were inoculated and grown at 37 °C in LB Lennox unless otherwise noted, and, when necessary, were supplemented with 20 mg/L chloramphenicol, 25 mg L^-1^ kanamycin, 10 mg L^-1^ tetracycline, or a combination thereof. Insertion-deletion alleles containing kanamycin resistance cassettes from the Keio collection (Baba et al., 2006) were moved to isogenic backgrounds using P1vir transduction (Silhavy et al., 1984). Clean deletions were constructed using the Flp recombinase-FRT system as described (Datsenko and Wanner, 2000). Chromosomal mutagenesis in *wzzE* was performed using a CRISPR-Cas9 system harboring a pCRISPR::protospacer plasmid in W3110 containing a heat shock-inducible λcI857 lysogen, as previously described (Pyne et al., 2015, Jiang et al., 2013). The pCas9 (Addgene #42876) and pCRISPR (Addgene #42975) plasmids were a kind gift from Professor Luciano Marraffini (Rockefeller University). Oligonucleotides used to generate the protospacer and repair templates used for mutagenesis are listed in **Table S3**. Prior to chromosomal mutagenesis, *metE*::Tn*10* was linked to *wzzE* (Nichols et al., 1998), allowing the mutations to be transferred into MG1655 via P1vir transduction. Strains carrying *metE*::Tn*10* were supplemented with 1µm cobalamin (vitamin B_12_) when grown in minimal media (Banerjee et al., 1989).

### Plasmids, cloning, and site directed mutagenesis

All primers used in this study are listed in **Table S4**. To subclone the p*wzzE* expression plasmid, chromosomal *wzzE* with an ochre stop codon was PCR-amplified using primers pZS21-wzzE_fwd and wzzE-pZS21_rev. The low copy number plasmid pZS21 (Lutz and Bujard, 1997) was amplified using primers pZS21_fwd and pZS21_rev. The plasmid was assembled using HiFi Assembly Master Mix (New England Biolabs) according to the manufacturer’s instructions. To construct p*wzzE-cat*, *wzzE* lacking a stop codon was amplified using primers pZS21-wzzE_fwd and wzzE-cat_rev. The chloramphenicol acetyltransferase gene (*cat*) was amplified from pBAD33 (Guzman et al., 1995) using primers wzzE-cat_fwd and cat-pZS21_rev. These fragments were assembled into pZS21, amplified as described above, using HiFi Assembly. To construct p*wzzE*-FLAG, p*wzzE* was amplified using primers p*wzzE*_x3F_Fwd and p*wzzE*_x3F_Rvr, which contained sequence homology to a glycine-serine-linked FLAG tag that incorporating a stop codon. The glycine-serine-linked FLAG tag was supplied as a gene-block fragment (**Table S3**) and assembled into the p*wzzE* plasmid via HiFi Assembly. To generate the p*wzzE-cat-*FLAG plasmid, *cat* was amplified using primers WzzE:Cmr Fp and WzzE:Cmr Rp and the p*wzzE*-FLAG plasmid backbone was amplified using Cmr: GS-x3F_FP and wzzE-cat_rev. The fragments were assembled using HiFi Assembly.

Site-direct mutations were generated using Q5 PCR mutagenesis (New England Biolabs). Briefly, non-overlapping primers containing the mutation of interest were used to amplify the indicated pZS21-sanA plasmids (pZS21-*wzzE*, pZS21-*wzzE-cat*, and pZS21-*wzzE*-FLAG). PCR products were digested with DpnI (New England Biolabs), phosphorylated using T4 polynucleotide kinase (New England Biolabs) and self-ligated with T4 DNA ligase (New England Biolabs). Ligations were transformed into Turbo cloning cells (New England Biolabs), and the mutations were confirmed by sequencing (Plasmidsaurs). Primers used all site-directed mutants are listed in **Table S4**. Modeling of the predicted mutant structures was performed using AlphaFold 3 (Abramson et al., 2024) with eight monomers of WzzE and one of WzyE. Relevant confidence scores are given in **Table S1** and **Figure S2**.

### Antibiotic Sensitivity

Efficiency of plating (EOP) assays were performed by plating 10-fold serial dilutions of overnight cultures onto LB plates containing the indicated antibiotics using a 48-pin tool. Plates were incubated overnight at 30 °C before imaging.

### Immunoblot Analysis

Immunoblots for WzzE-FLAG and ECA were conducted as previously described with minor modification (Mitchell et al., 2018, Morris and Mitchell, 2023). For analysis of WzzE-FLAG protein levels, samples were normalized to equal OD_600_ from overnight cultures, lysed by boiling for 5 minutes in sample loading buffer, and loaded on 12% SDS-PAGE gels.

Levels of WzzE-FLAG were quantitated by densitometry using ImageJ as has been previously described (Mitchell et al., 2018) on immunoblots with the lowest exposure at which WzzE bands could be detected. Samples for ECA analysis were prepared and immunoblotted as previously described with minor modification (Mitchell et al., 2018, Morris and Mitchell, 2023). To avoid ECA degradation, ultrapure water, commercial lysis reagents, commercial running buffer, and precast gels were used for ECA immunoblots. In addition, all western equipment was washed before and after ECA immunoblots with Extran 300 Detergent (Millipore Sigma) and rinsed 10 times with ultrapure water. To determine whether proteins affect the apparent chain length of ECA, we pre-treated samples with ProK. Samples were subjected to different pre-treatment conditions, varying both in the concentration of ProK and time of treatment (**Figure S5A**), based on protocols used in previous (Peters et al., 1985, Liu et al., 2020, Leo et al., 2020, Maczuga et al., 2022a) papers. All treatments were performed at 56 °C. Following ProK treatment, samples were boiled again for 10min, and loaded on 15% SDS-PAGE gels. The outcomes of each treatment condition and their respective effects on the resolution and clarity of the immunoblot, as well as effects on ECA levels are shown in (**Figure S5B**). Antibodies used for blotting were as follows: M2 αFLAG (Millipore Sigma, 1:50,000 dilution), αGroEL (Millipore Sigma, 1: 60,000 dilution), αECA (a gift from Professor Renato Morona, University of Adelaide, 1:30,000), rabbit α-mouse (Prometheus, 1:100,000 dilution), goat α-rabbit (Prometheus, 1:100,000 dilution), or αBamA (a gift from Professor Thomas Silhavy, Princeton University, 1:50,000 dilution).

### Quantification of Cyclic ECA

Quantification of ECA_CYC_ was performed as previously described, with minor modifications (Mitchell et al., 2018). Briefly, cells were grown in M63 glucose minimum media lacking nitrogen and supplemented with either 0.2% (^14^NH_4_)_2_SO_4_ or (^15^NH_4_)_2_SO_4_. ECA_CYC_ was purified as previously described. Sample analysis was performed using a Thermo Scientific Q Exactive Focus and liquid chromatography unit (UltiMate 3000 RS). Purified samples were dissolved in 0.1% formic acid and separated by injecting 20 µL of sample into the Synergi Hydro-RP (4.6 x 250 mm; 4.0 µm) column (Phenomenax). The mobile phase consisted of 0.1% Formic acid (eluent A) and methanol (eluent B). The flow rate was set at 800 µL/min with the following gradient: 0–5.0 min 5% B, 5.0-12.0 min 5-60% B, 12-14 min 60-90% B, 14-17 min 90% B, and returned to 5% B in 0.1 min and held for 5 min at 5% B. The Q Exactive Focus ESI source was operated in full MS (100-1000 m/z). The mass resolution was tuned to 70000 FWHM at m/z 200. The spray voltage was set to 3.3 kV in negative mode, and the sheath gas and auxiliary gas flow rates were set to 60 and 20 arbitrary units, respectively. The transfer capillary and auxiliary gas heater temperatures were held at 380 and 300 °C, respectively. The S-Lens RF level was set at 70 V. Exactive Series 2.11 /Xcalibur 4.2.47 software was used for data acquisition and processing. For relative quantification, the ratio of the areas of the ^14^N to ^15^N ECA_CYC_ peaks was calculated.

## Acknowledgements

Thank you to the members of the Mitchell Lab for helpful and productive discussions and for critical reading of our manuscript. We also thank Drs. Jolene Ramsey, Deborah Siegele, Jennifer Herman, and Ryland Young (Texas A&M University), as well as members of the Ramsey lab, for their insightful comments. The ECA antibody was a kind gift from Professor Renato Morana (University of Adelaide). The pCas9 and pCRISPR plasmids were a kind gift from Professor Luciano Marraffini (Rockefeller University) through the Addgene Repository. This work was supported by the National Institute of Allergy and Infectious Disease under award number R01-AI155915 (to A.M.M.).

