## Supplemental Figures and Tables for "Chain-length regulation by WzzE is necessary for, but genetically separable from, cyclic enterobacterial common antigen synthesis"

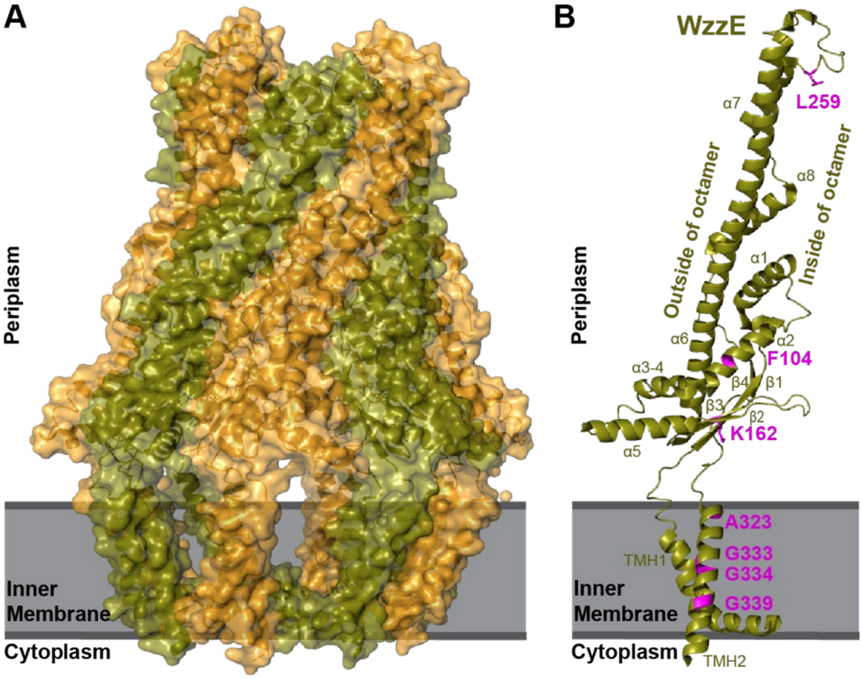


**Figure S1. Structure of WzzE. (A)** A cryoEM structure of the WzzE octamer (1) is shown with alternating monomers indicated by color. **(B)** The locations of residues changed in this study are called out in magenta on a WzzE monomer.

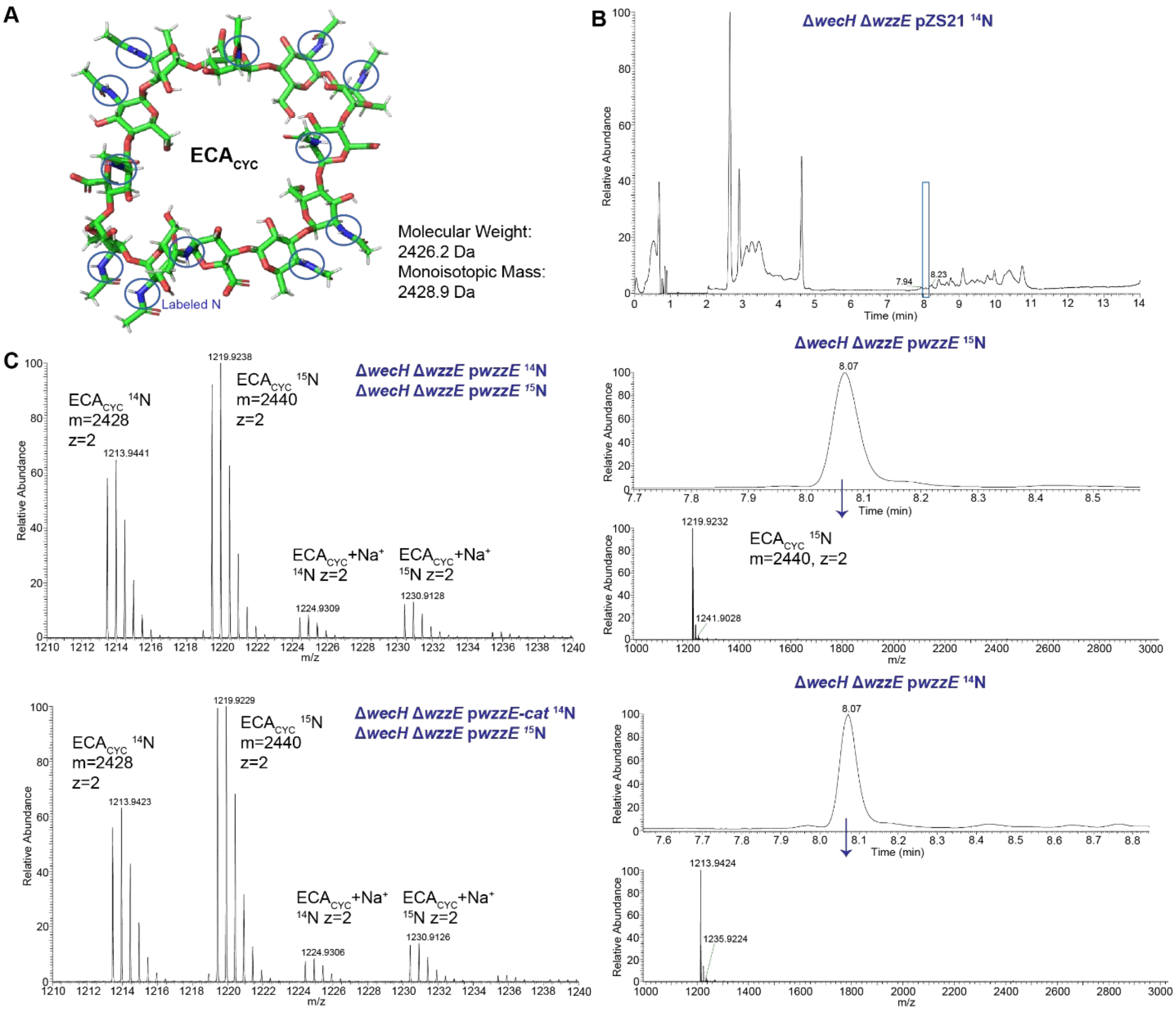


**Figure S2. Liquid chromatography-mass spectrometry (LC-MS) quantification of ECA$_{CYC}$. (A)** A crystal structure image of ECA$_{CYC}$ in its square conformation (2) is shown. The molecular weight and its monoisotopic mass, which takes into account the most common distributions of naturally occurring isotopes, are provided. The 12 nitrogen atoms (highlighted) will be made 1 Da heavier by growth with $^{15}$N, shifting the mass of ECA$_{CYC}$ by 12 Da. **(B)** LC and MS traces for individual samples of the indicated strains are shown. For the strain carrying the empty pZS21 vector, a blue box indicates the missing peak for ECA$_{CYC}$ (no MS trace is shown). **(C)** Representative MS traces are shown for samples from the indicated strains mixed before the start of ECA$_{CYC}$ purification. The ratio of the $^{14}$N and $^{15}$N peaks are nearly identical whether the sample grown in $^{14}$N carries the p*wzzE* or p*wzzE-cat* plasmid. The top image shown is the same data as in Figure 1E.

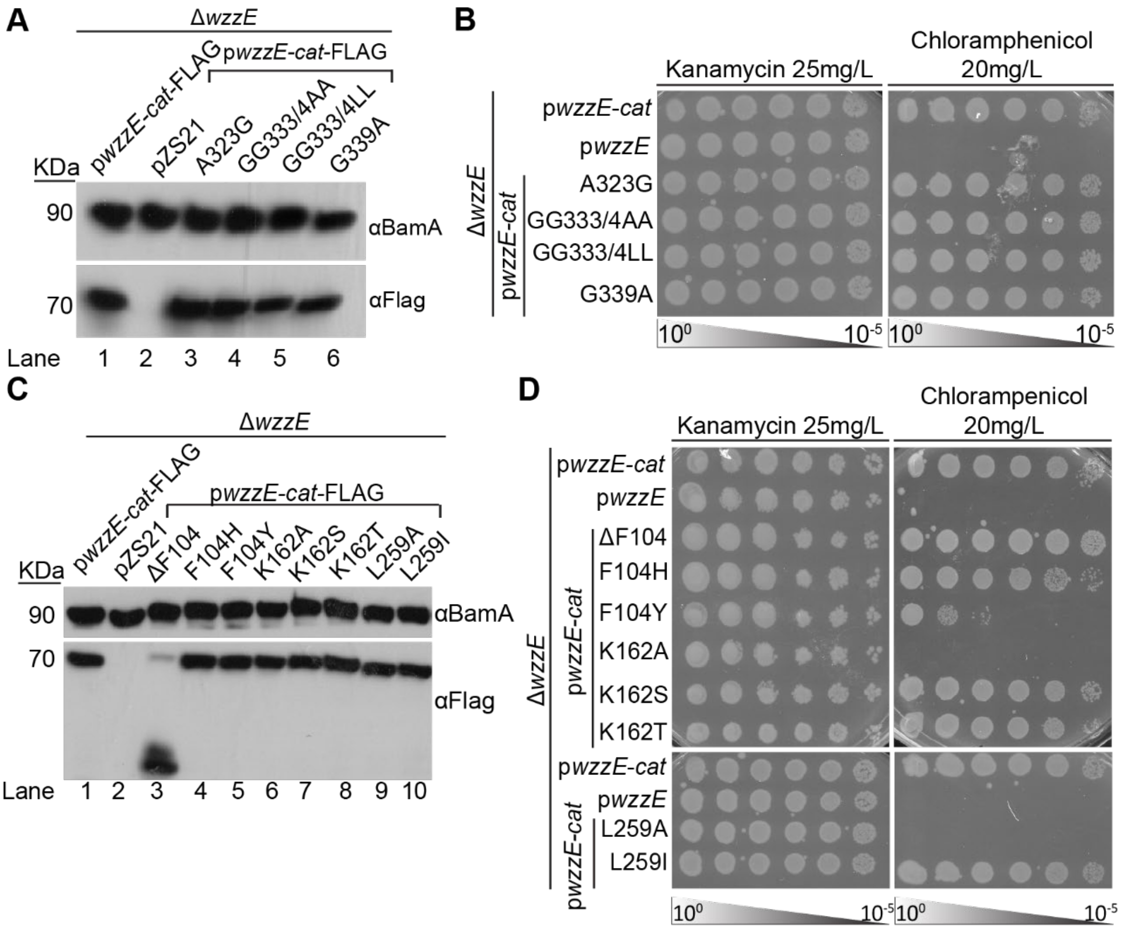


**Figure S3. Levels, stability, and multimerization of *wzzE* mutants. (A)** The effect of the indicated mutations to the region of *wzzE* encoding TMH2 made in the p*wzzE-cat*-FLAG background was assayed by immunoblotting. BamA serves as a loading control. All mutants appear to be equally stable. **(B)** EOPs are shown assaying the stability and multimerization of WzzE using the proxy of chloramphenicol resistance. Mutations are expressed from the p*wzzE-cat* construct. All mutants retain equal chloramphenicol resistance. **(C)** The effect of the indicated mutations to the region of *wzzE* encoding the periplasmic domain made in the p*wzzE-cat*-FLAG background was assayed by immunoblotting. BamA serves as a loading control. All mutants except $wzzE_{\Delta F104}$ appear to be equally stable. The C-terminal tags are largely cleaved from $WzzE^{\Delta F104}$ making its stability unclear. **(D)** EOPs are shown assaying the stability and multimerization of WzzE using the proxy of chloramphenicol resistance in mutants expressed from the p*wzzE-cat* construct. Several mutants show decreased chloramphenicol resistance, including F104Y, K162A, and L259A.

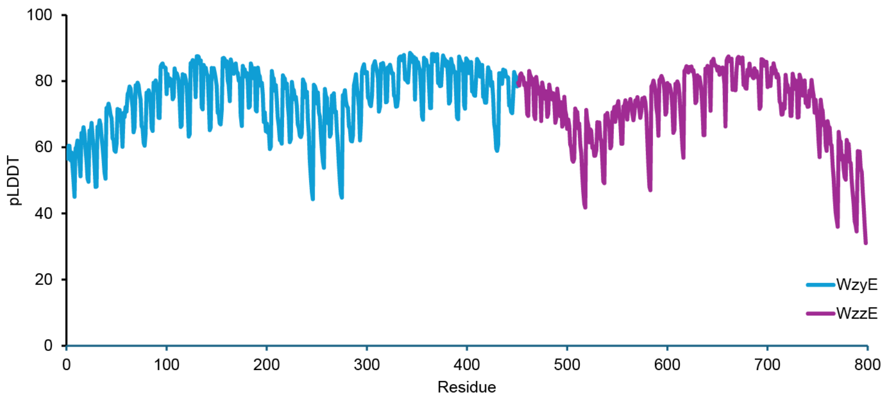


**Figure S4. Confidence scores for WzzE-WzyE complex structural prediction.** Predicted local distance difference test (pLDDT) scores are shown for WzyE and the first WzzE monomer from an AlphaFold 3 (3) prediction of the complex of wild-type WzzE(8) with WzyE. Distributions were similar for the predicted structures with WzzE mutants.

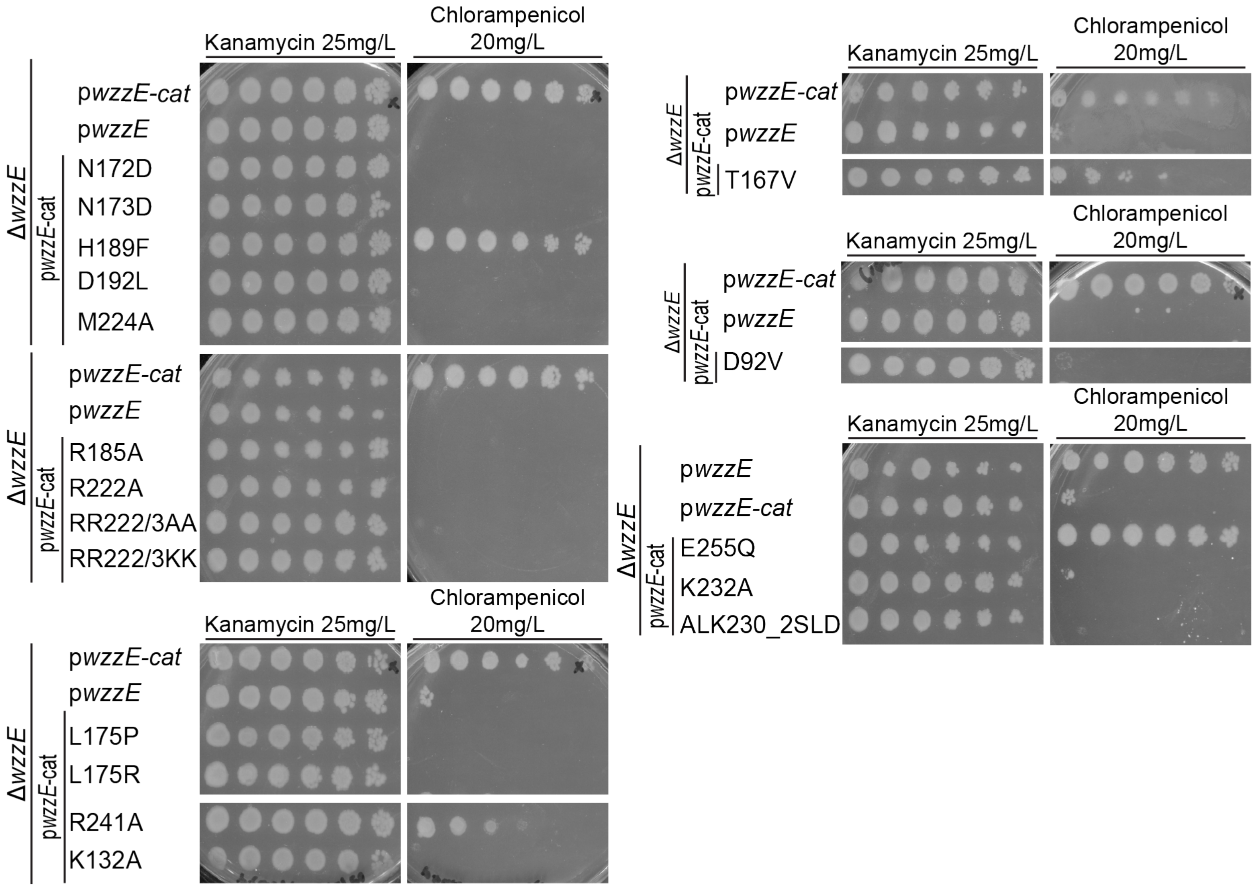


**Figure S5. Changes to the periplasmic domain of WzzE are prone to induce instability and/or affect multimerization.** Strains with the indicated plasmids were assayed for chloramphenicol resistance as a measure of their stability and ability to multimerize. These periplasmic mutations led to loss of chloramphenicol resistance indicating that the mutant proteins were unstable or unable to multimerize.

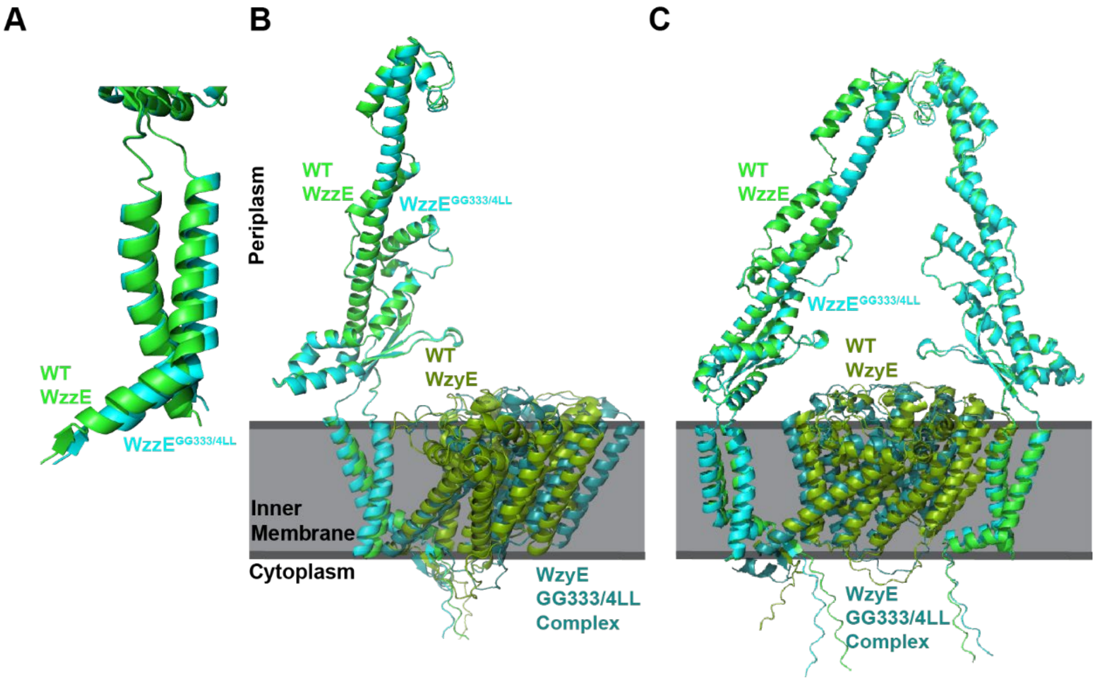


**Figure S6. WzzE-WzyE interactions may shift in the WzzE$^{GG333/4LL}$ mutant.** Models of the WzzE(8)-WzyE complex were constructed with Alphafold3 for wild-type WzzE and the WzzE$^{GG333/4LL}$ mutant and the WzzE monomers most closely contacting WzyE were aligned. **(A)** The TMHs of WzzE are closer together in the wild-type model than in the WzzE$^{GG333/4LL}$ model. **(B-C)** The aligned WzzE monomers are shown with WzyE (B) or the aligned monomers and the opposite monomers (C). The predicted angle of WzyE shifts in the WzzE$^{GG333/4LL}$ model compared with the wild type.

**A**

| Treatment | Protinase K concentration | Incubation time |
|---|---|---|
| T1 | 1.6 mg/ml | overnight |
| T2 | 0.02 mg/ml | overnight |
| T3 | 0.25 mg/ml | overnight |
| T4 | 2.5 mg/ml | 2 hours |
| T5 | 2.5 mg/ml | 1 hour |

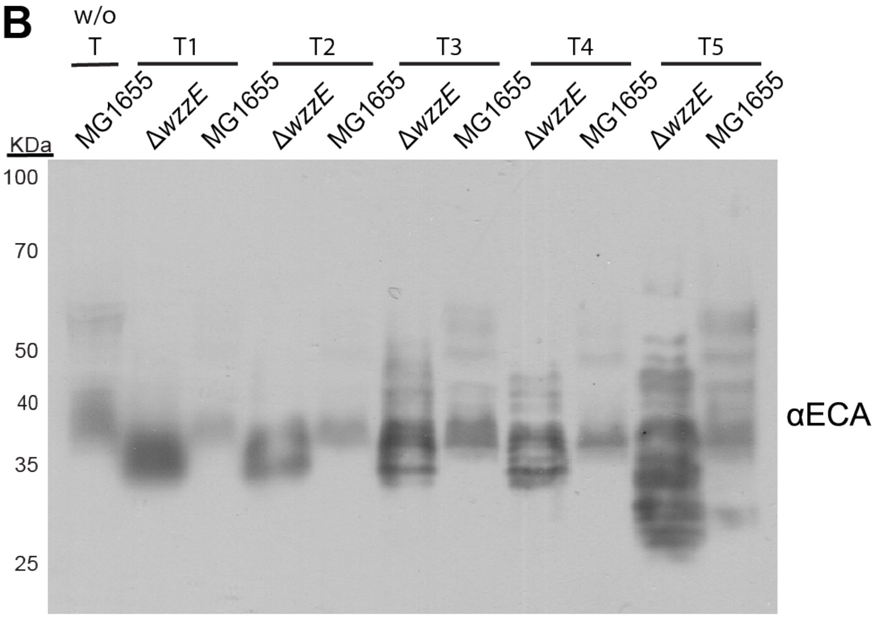


**Figure S7. The effect of proteinase K treatment on ECA immunoblots. (A)** A summary of proteinase K treatment conditions prior to ECA immunoblotting is shown. Five treatments (T1–T5) conditions were chosen from or modified from literature (4-8) and varied in proteinase K concentration added and time of digestion at 56 °C. **(B)** Immunoblot analysis of ECA following treatment. Whole-cell lysates from wild-type *and* Δ*wzzE* were subjected to the treatments described above and probed with anti-ECA antibodies. For each treatment, the wild-type and ΔwzzE samples were processed and run in parallel to compare the impact of proteinase K treatment on the banding pattern and ECA stability. The far-left lane ("w/o T") represents an untreated wild-type control, harvested via our normal protocol. Longer treatments resulted in a lower signal ECA signal, likely because of ECA degradation. However, a short incubation with high amounts of proteinase K resulted in more even banding without significant loss of ECA.

**SUPPLEMENTAL TABLES**

**Table S1: Confidence scores for WzzE-WzyE structural predictions**

| Model | pTM[1] | ipTM[2] |
|---|---|---|
| **WzzE(8)-WzyE** | 0.75 | 0.74 |
| **WzzE$^{GG333/4LL}$-WzyE** | 0.74 | 0.73 |
| **WzzE$^{F104\Delta}$-WzyE** | 0.73 | 0.71 |
| **WzzE$^{F104H}$-WzyE** | 0.75 | 0.74 |
| **WzzE$^{F104Y}$-WzyE** | 0.75 | 0.74 |
| **WzzE(8) only** | 0.80 | 0.78 |

[1] Predicted modeling template score, ranging from 0-1, indicating the confidence in the overall structure of the complex.
[2] Interface predicted template modeling score, ranging from 0-1, indicating the confidence that relative position of the complex subunits is correct.

**Table S2: Strains used in this study**

| Strain | Genotype | Reference |
|---|---|---|
| MG1655 | K-12 $F^{-}$ $\lambda^{-}$ *rph-1* | (9) |
| AM369 | MG1655 Δ*wzzE* Δ*yhdP* | (10) |
| AM365 | MG1655 Δ*wzzE* | (10) |
| AM182 | MG1655 Δ*yhdP* | (10) |
| AM1113 | MG1655 Δ*wzzE* Δ*yhdP* pZS21 | This study |
| AM1114 | MG1655 Δ*wzzE* Δ*yhdP* p*wzzE* | This study |
| AM1116 | MG1655 Δ*wzzE* Δ*yhdP* p*wzzE-cat* | This study |
| JC118 | MG1655 Δ*wzzE* Δ*yhdP* p*wzzE-FLAG* | This study |
| JC205 | MG1655 Δ*wzzE* pZS21 | This study |
| JC206 | MG1655 Δ*wzzE* p*wzzE* | This study |
| JC207 | MG1655 Δ*wzzE* p*wzzE*-FLAG | This study |
| JC155 | MG1655 Δ*wzzE* Δ*wecH* | This study |
| JC110 | MG1655 Δ*wecH* | This study |
| JC159 | MG1655 Δ*wzzE* Δ*wecH* pZS21 | This study |
| JC160 | MG1655 Δ*wzzE* ΔwecH p*wzzE* | This study |
| AM335 | MG1655 Δ*wecA* | This study |
| JC225 | MG1655 Δ*wzzE* p$wzzE_{A323G}$ | This study |
| JC226 | MG1655 Δ*wzzE* p$wzzE_{GG333/4AA}$ | This study |
| JC227 | MG1655 Δ*wzzE* p$wzzE_{GG333/4LL}$ | This study |
| JC228 | MG1655 Δ*wzzE* p$wzzE_{G339A}$ | This study |
| JC35 | MG1655 Δ*wzzE* p$wzzE_{GG333/4AA}$-*cat* | This study |
| JC36 | MG1655 Δ*wzzE* p$wzzE_{A323G}$-*cat* | This study |
| JC40 | MG1655 Δ*wzzE* p$wzzE_{GG333/4LL}$-*cat* | This study |
| JC41 | MG1655 Δ*wzzE* p$wzzE_{G339A}$-*cat* | This study |
| JC210 | MG1655 Δ*wzzE* p$wzzE_{A323G}$-FLAG | This study |
| JC211 | MG1655 Δ*wzzE* p$wzzE_{GG333/4AA}$-FLAG | This study |
| JC212 | MG1655 Δ*wzzE* p$wzzE_{GG333/4LL}$-FLAG | This study |
| JC213 | MG1655 Δ*wzzE* p$wzzE_{G339A}$-FLAG | This study |
| JC381 | MG1655 Δ*wzzE* p$wzzE_{A323G}$-*cat*-FLAG | This study |
| JC382 | MG1655 Δ*wzzE* p$wzzE_{GG333/4AA}$-*cat*-FLAG | This study |
| JC383 | MG1655 Δ*wzzE* p$wzzE_{GG333/4LL}$-*cat*-FLAG | This study |
| JC385 | MG1655 Δ*wzzE* p$wzzE_{G339A}$-*cat*-FLAG | This study |
| JC163 | MG1655 Δ*wzzE* Δ*wecH* p$wzzE_{GG333/4LL}$ | This study |
| JC167 | MG1655 Δ*wzzE* Δ*wecH* p$wzzE_{A323G}$ | This study |
| JC168 | MG1655 Δ*wzzE* Δ*wecH* p$wzzE_{GG333/4AA}$ | This study |
| JC170 | MG1655 Δ*wzzE* Δ*wecH* p$wzzE_{G339A}$ | This study |
| JC44 | MG1655 Δ*wzzE* p$wzzE_{\Delta F104}$-*cat* | This study |
| JC45 | MG1655 Δ*wzzE* p$wzzE_{F104H}$-*cat* | This study |
| JC46 | MG1655 Δ*wzzE* p$wzzE_{F104Y}$-*cat* | This study |
| JC47 | MG1655 Δ*wzzE* p$wzzE_{L259A}$-*cat* | This study |

| Strain | Genotype | Reference |
|---|---|---|
| JC48 | MG1655 Δ*wzzE* p$wzzE_{\mathrm{L259I}}$-*cat* | This study |
| JC50 | MG1655 Δ*wzzE* p$wzzE_{\mathrm{K162A}}$-*cat* | This study |
| JC51 | MG1655 Δ*wzzE* p$wzzE_{\mathrm{K162S}}$-*cat* | This study |
| JC52 | MG1655 Δ*wzzE* p$wzzE_{\mathrm{K162T}}$-*cat* | This study |
| JC208 | MG1655 Δ*wzzE* p$wzzE_{\Delta \mathrm{F104}}$-FLAG | This study |
| JC209 | MG1655 Δ*wzzE* p$wzzE_{\mathrm{F104H}}$-FLAG | This study |
| JC214 | MG1655 Δ*wzzE* p$wzzE_{\mathrm{F104Y}}$-FLAG | This study |
| JC215 | MG1655 Δ*wzzE* p$wzzE_{\mathrm{K162A}}$-FLAG | This study |
| JC216 | MG1655 Δ*wzzE* p$wzzE_{\mathrm{K162S}}$-FLAG | This study |
| JC217 | MG1655 Δ*wzzE* p$wzzE_{\mathrm{K162T}}$-FLAG | This study |
| JC218 | MG1655 Δ*wzzE* p$wzzE_{\mathrm{L259A}}$-FLAG | This study |
| JC219 | MG1655 Δ*wzzE* p$wzzE_{\mathrm{L259I}}$-FLAG | This study |
| JC226 | MG1655 Δ*wzzE* p$wzzE_{\Delta \mathrm{F104}}$ | This study |
| JC227 | MG1655 Δ*wzzE* p$wzzE_{\mathrm{F104Y}}$ | This study |
| JC228 | MG1655 Δ*wzzE* p$wzzE_{\mathrm{F104H}}$ | This study |
| JC221 | MG1655 Δ*wzzE* p$wzzE_{\mathrm{K162A}}$ | This study |
| JC220 | MG1655 Δ*wzzE* p$wzzE_{\mathrm{K162S}}$ | This study |
| JC231 | MG1655 Δ*wzzE* p$wzzE_{\mathrm{K162T}}$ | This study |
| JC229 | MG1655 Δ*wzzE* p$wzzE_{\mathrm{L259A}}$ | This study |
| JC230 | MG1655 Δ*wzzE* p$wzzE_{\mathrm{L259I}}$ | This study |
| JC376 | MG1655 Δ*wzzE* p*wzzE*-*cat*-FLAG | This study |
| JC373 | MG1655 Δ*wzzE* p$wzzE_{\Delta \mathrm{F104}}$-*cat*-FLAG | This study |
| JC374 | MG1655 Δ*wzzE* p$wzzE_{\mathrm{F104H}}$-*cat*-FLAG | This study |
| JC375 | MG1655 Δ*wzzE* p$wzzE_{\mathrm{F104Y}}$-*cat*-FLAG | This study |
| JC380 | MG1655 Δ*wzzE* p$wzzE_{\mathrm{L259A}}$-*cat*-FLAG | This study |
| JC384 | MG1655 Δ*wzzE* p$wzzE_{\mathrm{L259I}}$-*cat*-FLAG | This study |
| JC377 | MG1655 Δ*wzzE* p$wzzE_{\mathrm{K162A}}$-*cat*-FLAG | This study |
| JC378 | MG1655 Δ*wzzE* p$wzzE_{\mathrm{K162S}}$-*cat*-FLAG | This study |
| JC379 | MG1655 Δ*wzzE* p$wzzE_{\mathrm{K162T}}$-*cat*-FLAG | This study |
| JC171 | MG1655 Δ*wzzE* Δ*wecH* p$wzzE_{\Delta \mathrm{F104}}$ | This study |
| JC172 | MG1655 Δ*wzzE* Δ*wecH* p$wzzE_{\mathrm{F104H}}$ | This study |
| JC173 | MG1655 Δ*wzzE* Δ*wecH* p$wzzE_{\mathrm{F104Y}}$ | This study |
| JC174 | MG1655 Δ*wzzE* Δ*wecH* p$wzzE_{\mathrm{K162A}}$ | This study |
| JC175 | MG1655 Δ*wzzE* Δ*wecH* p$wzzE_{\mathrm{K162S}}$ | This study |
| JC176 | MG1655 Δ*wzzE* Δ*wecH* p$wzzE_{\mathrm{L259A}}$ | This study |
| JC177 | MG1655 Δ*wzzE* Δ*wecH* p$wzzE_{\mathrm{K162T}}$ | This study |
| JC178 | MG1655 Δ*wzzE* Δ*wecH* p$wzzE_{\mathrm{L259I}}$ | This study |
| JC308 | MG1655 $wzzE_{\Delta F104}$ Δ*wecH* | This study |
| JC312 | MG1655 $wzzE_{GG333/4LL}$ | This study |
| JC288 | MG1655 $wzzE_{GG333/4LL}$ Δ*wecH* | This study |
| JC321 | MG1655 $wzzE_{F104Y}$ Δ*wecH* | This study |

| **Strain** | **Genotype** | **Reference** |
| --- | --- | --- |
| JC322 | MG1655 *wzzE*$_{F104H}$ Δ*wecH* | This study |
| JC324 | MG1655 *wzzE*$_{F104H}$ | This study |
| JC325 | MG1655 *wzzE*$_{F104Y}$ | This study |
| JC326 | MG1655 *wzzE*$_{\Delta F104}$ | This study |
| JC340 | MG1655 Δ*wzzB* | This study |
| JC341 | MG1655 Δ*fepE* | This study |
| JC342 | MG1655 Δ*wzzE* Δ*wzzB:*Kan | This study |
| JC343 | MG1655 Δ*wzzE* Δ*fepE:*Kan | This study |
| JC351 | MG1655 Δ*wzzE* Δ*wzzB* Δ*fepE* | This study |
| JC53 | MG1655 Δ*wzzE* p*wzzE*$_{N172D}$-*cat* | This study |
| JC54 | MG1655 Δ*wzzE* p*wzzE*$_{N173D}$-*cat* | This study |
| JC55 | MG1655 Δ*wzzE* p*wzzE*$_{H189F}$-*cat* | This study |
| JC56 | MG1655 Δ*wzzE* p*wzzE*$_{D192L}$-*cat* | This study |
| JC60 | MG1655 Δ*wzzE* p*wzzE*$_{R185A}$-*cat* | This study |
| JC61 | MG1655 Δ*wzzE* p*wzzE*$_{R222A}$-*cat* | This study |
| JC62 | MG1655 Δ*wzzE* p*wzzE*$_{RR222/3AA}$-*cat* | This study |
| JC63 | MG1655 Δ*wzzE* p*wzzE*$_{RR222/3KK}$-*cat* | This study |
| JC28 | MG1655 Δ*wzzE* p*wzzE*$_{L175P}$-*cat* | This study |
| JC29 | MG1655 Δ*wzzE* p*wzzE*$_{L175R}$-*cat* | This study |
| JC30 | MG1655 Δ*wzzE* p*wzzE*$_{D92V}$-*cat* | This study |
| JC31 | MG1655 Δ*wzzE* p*wzzE*$_{K132A}$-*cat* | This study |
| JC64 | MG1655 Δ*wzzE* p*wzzE*$_{M224A}$-*cat* | This study |
| JC65 | MG1655 Δ*wzzE* p*wzzE*$_{T167V}$-*cat* | This study |
| JC80 | MG1655 Δ*wzzE* p*wzzE*$_{E225Q}$-*cat* | This study |
| JC81 | MG1655 Δ*wzzE* p*wzzE*$_{K232A}$-*cat* | This study |
| JC82 | MG1655 Δ*wzzE* p*wzzE*$_{ALK230\_2SLD}$-*cat* | This study |

**Table S3: CRISPR spacer oligos and repair templates**

| Oligo/Geneblock | Sequence 5'→3' |
|---|---|
| **ΔF104 Repair Template Gene Block** | atgACACAACCAATGCCTGGGAAACCGGCCGAAGACGCT<br>GAAAATGAACTGGATATTCGTGGGTTGTTTCGTACCTTGTG<br>GGCTGGGAAGCTATGGATTATTGGCATGGGGCTGGCGTT<br>TGCGTTAATCGCGCTGGCGTATACGTTTTTTGCTCGTCAG<br>GAGTGGAGCTCGACGGCGATTACCGATCGTCCAACGGT<br>GAATATGCTGGGGGGATATTACTCGCAGCAGCAATTTTTG<br>CGTAACCTGGATGTCCGTTCAAACATGGCTTCTGCCGAC<br>CAACCATCGGTCATGGACGAAGCCTACAAAGAGGTTATG<br>CAGCTGGCCAGTTGGGATACCCGCAGAGAGTTCTGGCT<br>GCAAACCGACTATTACAAACAGCGGATGGTGGGCAACA<br>GCAAAGCCGATGCGGCGTTGCTGGATGAAATGATTAACA<br>ACATCCAGTTTATCCCCGGAGACTTTACCCGCGCGGTCA<br>ATGACAGCGTGAAGCTTATTGCCGAAACCGCGCCTGAC<br>GCTAATAACCTGTTACGTCAGTATGTTGCTTTTGCCAGCC<br>AGCGTGCAGCCAGCCATCTGAATGATGAGCTGAAAGGC<br>GCATGGGCGGCGCGTACCATCCAGATGAAAGCTCAGGT<br>GAAGCGTCAGGAAGAGGTGGCGAAAGCCATCTACGACC<br>GCCGGATGAACAGCATTGAGCAGGCGCTGAAAATTGCT<br>GAGCAGCATAATATTTCGCGCAGTGCGACAGATGTACCT<br>GCCGAGGAATTA |
| **F104H Repair Template Gene Block** | atgACACAACCAATGCCTGGGAAACCGGCCGAAGACGCT<br>GAAAATGAACTGGATATTCGTGGGTTGTTTCGTACCTTGTG<br>GGCTGGGAAGCTATGGATTATTGGCATGGGGCTGGCGTT<br>TGCGTTAATCGCGCTGGCGTATACGTTTTTTGCTCGTCAG<br>GAGTGGAGCTCGACGGCGATTACCGATCGTCCAACGGT<br>GAATATGCTGGGGGGATATTACTCGCAGCAGCAATTTTTG<br>CGTAACCTGGATGTCCGTTCAAACATGGCTTCTGCCGAC<br>CAACCATCGGTCATGGACGAAGCCTACAAAGAGCACGTT<br>ATGCAGCTGGCCAGTTGGGATACCCGCAGAGAGTTCTG<br>GCTGCAAACCGACTATTACAAACAGCGGATGGTGGGCAA<br>CAGCAAAGCCGATGCGGCGTTGCTGGATGAAATGATTAA<br>CAACATCCAGTTTATCCCCGGAGACTTTACCCGCGCGGT<br>CAATGACAGCGTGAAGCTTATTGCCGAAACCGCGCCTGA<br>CGCTAATAACCTGTTACGTCAGTATGTTGCTTTTGCCAGC<br>CAGCGTGCAGCCAGCCATCTGAATGATGAGCTGAAAGG<br>CGCATGGGCGGCGCGTACCATCCAGATGAAAGCTCAGG<br>TGAAGCGTCAGGAAGAGGTGGCGAAAGCCATCTACGAC<br>CGCCGGATGAACAGCATTGAGCAGGCGCTGAAAATTGCT<br>GAGCAGCATAATATTTCGCGCAGTGCGACAGATGTACCT<br>GCCGAGGAA |
| **F104Y Repair Template Gene Block** | atgACACAACCAATGCCTGGGAAACCGGCCGAAGACGCT<br>GAAAATGAACTGGATATTCGTGGGTTGTTTCGTACCTTGTG |

| Oligo/Geneblock | Sequence 5'→3' |
|---|---|
| | GGCTGGGAAGCTATGGATTATTGGCATGGGGCTGGCGTT<br>TGCGTTAATCGCGCTGGCGTATACGTTTTTTGCTCGTCAG<br>GAGTGGAGCTCGACGGCGATTACCGATCGTCCAACGGT<br>GAATATGCTGGGGGGATATTACTCGCAGCAGCAATTTTTG<br>CGTAACCTGGATGTCCGTTCAAACATGGCTTCTGCCGAC<br>CAACCATCGGTCATGGACGAAGCCTACAAAGAGTACGTT<br>ATGCAGCTGGCCAGTTGGGATACCCGCAGAGAGTTCTG<br>GCTGCAAACCGACTATTACAAACAGCGGATGGTGGGCAA<br>CAGCAAAGCCGATGCGGCGTTGCTGGATGAAATGATTAA<br>CAACATCCAGTTTATCCCCGGAGACTTTACCCGCGCGGT<br>CAATGACAGCGTGAAGCTTATTGCCGAAACCGCGCCTGA<br>CGCTAATAACCTGTTACGTCAGTATGTTGCTTTTGCCAGC<br>CAGCGTGCAGCCAGCCATCTGAATGATGAGCTGAAAGG<br>CGCATGGGCGGCGCGTACCATCCAGATGAAAGCTCAGG<br>TGAAGCGTCAGGAAGAGGTGGCGAAAGCCATCTACGAC<br>CGCCGGATGAACAGCATTGAGCAGGCGCTGAAAATTGCT<br>GAGCAGCATAATATTTCGCGCAGTGCGACAGATGTACCT<br>GCCGAGGAA |
| **GG333/4LL Repair Template Gene Block** | ATGACACAACCAATGCCTGGGAAACCGGCCGAAGACGC<br>TGAAAATGAACTGGATATTCGTGGGTTGTTTCGTACCTTGT<br>GGGCTGGGAAGCTATGGATTATTGGCATGGGGCTGGCGT<br>TTGCGTTAATCGCGCTGGCGTATACGTTTTTTGCTCGTCA<br>GGAGTGGAGCTCGACGGCGATTACCGATCGTCCAACGG<br>TGAATATGCTGGGGGGATATTACTCGCAGCAGCAATTTTT<br>GCGTAACCTGGATGTCCGTTCAAACATGGCTTCTGCCGA<br>CCAACCATCGGTCATGGACGAAGCCTATAAAGAGTTTGTT<br>ATGCAACTGGCCTCGTGGGATACCCGCAGAGAGTTCTG<br>GCTGCAAACCGACTATTACAAACAGCGGATGGTGGGCAA<br>CAGCAAAGCCGATGCGGCGTTGCTGGATGAAATGATTAA<br>CAACATCCAGTTTATCCCCGGAGACTTTACCCGCGCGGT<br>CAATGACAGCGTGAAGCTTATTGCCGAAACCGCGCCTGA<br>CGCTAATAACCTGTTACGTCAGTATGTTGCTTTTGCCAGC<br>CAGCGTGCAGCCAGCCATCTGAATGATGAGCTGAAAGG<br>CGCATGGGCGGCGCGTACCATCCAGATGAAAGCTCAGG<br>TGAAGCGTCAGGAAGAGGTGGCGAAAGCCATCTACGAC<br>CGCCGGATGAACAGCATTGAGCAGGCGCTGAAAATTGCT<br>GAGCAGCATAATATTTCGCGCAGTGCGACAGATGTACCT<br>GCCGAGGAATTACCTGATTCAGAAATGTTCCTGCTTGGG<br>CGTCCAATGCTTCAGGCTCGACTGGAAAATTTACAGGCC<br>GTCGGTCCGGCCTTTGATCTCGACTATGATCAGAATCGG<br>GCCATGTTAAACACCCTGAATGTTGGTCCAACCCTGGAT<br>CCGCGTTTTCAGACCTATCGCTATTTGCGTACGCCGGAA<br>GAACCGGTAAAACGCGATAGCCCACGTCGTGCCTTCCT |

| Oligo/Geneblock | Sequence 5'→3' |
|---|---|
| | GATGATTATGTGGGGCATTGTCcttcttCTGATCGGGGCTG<br>GTGTCGCATTAACCCGCCGTTGCTCGAAATAG |
| **F104 Protospacer Fwd** | AAACTATAAAGAGTTTGTTATGCAACTGGCCTCGG |
| **F104 Protospacer Rvr** | AAAACCGAGGCCAGTTGCATAACAAACTCTTTATA |
| **GG333/4LL Protopacer Fwd** | AAACTTCCTGATGATTATGTGGGGCATTGTCGGGG |
| **GG333/4LL protospacer Rvr** | AAAACCCCGACAATGCCCCACATAATCATCAGGAA |
| **wzzE_x3F Gene Block** | GGTGTCGCATTCCGCCGTTGCTCGAAAGGAGGCAGCGG<br>TGGGAGTGGCGGAGACTACAAGGACCACGACGGTGACT<br>ACAAGGACCACGACATCGACTACAAGGACGACGACGAC<br>AAGTGRAAAGTGGATCCGTCGACCTGCAGCCA |

**Table S4: Primers used in this study**

| Primer | Sequence 5' → 3' |
|---|---|
| pZS21_fwd | agtggatccgtcgacctg |
| pZS21_rev | catggtacctttctttaatgaattcgg |
| pZS21-wzzE_fwd | cattaaagaaaggtaccatggcgtagaaatcgtggtgg |
| wzzE-pZS21_rev | tgcaggtcgacggatccactttatttcgagcaacggcgg |
| wzzE-cat_rev | ttttctccattttcgagcaacggcgggt |
| wzzE-cat_fwd | ttgctcgaaaatggagaaaaaaatcactggatataccaccgttgatatat ccc |
| cat-pZS21_rev | tgcaggtcgacggatccactttacgccccgccctgcca |
| pZS21-wzzE_fwd ΔF104 | GTTATGCAACTGGCCTCG |
| pZS21-wzzE_Rvr ΔF104 | CTCTTTATAGGCTTCGTCC |
| pZS21-wzzE_fwd F104H | CTATAAAGAGcacGTTATGCAACTGGCCTC |
| pZS21-wzzE_Rvr F104H | GCTTCGTCCATGACCGAT |
| pZS21-wzzE_fwd F104Y | CTATAAAGAGtatGTTATGCAACTGG |
| pZS21-wzzE_Rvr F104Y | GCTTCGTCCATGACCGAT |
| pZS21-wzzE_fwd K162A | TGACAGCGTGgctCTTATTGCCGAAACCG |
| pZS21-wzzE_Rvr K162A | TTGACCGCGCGGGTAAAG |
| pZS21-wzzE_fwd K162S | TGACAGCGTGtccCTTATTGCCGAAACCG |
| pZS21-wzzE_Rvr K162S | TTGACCGCGCGGGTAAAG |
| pZS21-wzzE_fwd K162T | TGACAGCGTGactCTTATTGCCGAAACCGC |
| pZS21-wzzE_Rvr K162T | TTGACCGCGCGGGTAAAG |
| pZS21-wzzE_fwd L259A | AATGTTCCTGgctGGGCGTCCAATG |
| pZS21-wzzE_Rvr L259A | TCTGAATCAGGTAATTCCTC |
| pZS21-wzzE_Rvr L259I | TCTGAATCAGGTAATTCCTC |
| pZS21-wzzE_Fwd L259I | AATGTTCCTGattGGGCGTCCAA |
| pZS21-wzzE_fwd N172D | GCCTGACGCTgacAACCTGTTAC |
| pZS21-wzzE_Rvr N172D | GCGGTTTCGGCAATAAGC |
| pZS21-wzzE_fwd N173D | TGACGCTAATgatCTGTTACGTCAGTATGTTG |
| pZS21-wzzE_Rvr N173D | GCGGTTTCGGCAATAAGC |
| pZS21-wzzE_fwd H189F | TGCAGCCAGCttcCTGAATGATGAGCTGAAAG |
| pZS21-wzzE_Rvr H189F | CGCTGGCTGGCAAAAGCA |
| pZS21-wzzE_fwd D192L | CCATCTGAATcttGAGCTGAAAGGCGCATGG |
| pZS21-wzzE_Rvr D192L | CTGGCTGCACGCTGGCTG |
| pZS21-wzzE_fwd M224A | CGACCGCCGGgccAACAGCATTG |
| pZS21-wzzE_Rvr M224A | TAGATGGCTTTCGCCACC |
| pZS21-wzzE_fwd R185A | TGCCAGCCAGgccGCAGCCAGCC |
| pZS21-wzzE_Rvr R185A | AAAGCAACATACTGACGTAACAG |
| pZS21-wzzE_fwd RR222/3AA | CATCTACGACgccgccATGAACAGCATTGAG |
| pZS21-wzzE_Rvr RR222/3AA | GCTTTCGCCACCTCTTCC |
| pZS21-wzzE_fwd RR222/3KK | CATCTACGACaaaaaaATGAACAGCATTGAGCAG |
| pZS21-wzzE_Rvr RR222/3KK | GCTTTCGCCACCTCTTCC |

| Primer | Sequence 5' → 3' |
|---|---|
| pZS21-wzzE_fwd L175P | TAATAACCTGcctCGTCAGTATGTTGCTTTTGCCAGCC |
| pZS21-wzzE_Rvr L175P | GCGTCAGGCGCGGTTTCG |
| pZS21-wzzE_fwd L175R | TAATAACCTGcgtCGTCAGTATGTTGCTTTTGCCAGCC |
| pZS21-wzzE_Rvr L175R | GCGTCAGGCGCGGTTTCG |
| pZS21-wzzE_fwd R241A | TAATATTTCGgccAGTGCGACAGATG |
| pZS21-wzzE_Rvr R241A | TGCTGCTCAGCAATTTTC |
| pZS21-wzzE_fwd R222A | CATCTACGACgccCGGATGAACAGCATTG |
| pZS21-wzzE_Rvr R222A | GCTTTCGCCACCTCTTCC |
| pZS21-wzzE_fwd T167V | TATTGCCGAAgttGCGCCTGACG |
| pZS21-wzzE_Rvr T167V | AGCTTCACGCTGTCATTG |
| pZS21-wzzE_fwd D92V | GGCTTCTGCCgttCAACCATCGG |
| pZS21-wzzE_Rvr D92V | ATGTTTGAACGGACATCC |
| pZS21-wzzE_fwd E255Q | ACCTGATTCAcaaATGTTCCTGCTTG |
| pZS21-wzzE_Rvr E255Q | AATTCCTCGGCAGGTACA |
| pZS21-wzzE_fwd K232A | GCAGGCGCTGgccATTGCTGAGC |
| pZS21-wzzE_Rvr K232A | TCAATGCTGTTCATCCG |
| pZS21-wzzE_fwd ALK230_2SLD | agacATTGCTGAGCAGCATAATATTTC |
| pZS21-wzzE_Rvr ALK230_2SLD | aaggaCTGCTCAATGCTGTTCATC |
| pZS21-wzzE_fwd A323G | CCCACGTCGTggtTTCCTGATGATTATG |
| pZS21-wzzE_Rvr A323G | CTATCGCGTTTTACCGGT |
| pZS21-wzzE_fwd GG333/4AA | GGGCATTGTCgctgctCTGATCGGGG |
| pZS21-wzzE_Rvr GG333/4AA | CACATAATCATCAGGAAGG |
| pZS21-wzzE_fwd GG333/4LL | GGGCATTGTCcttcttCTGATCGGGG |
| pZS21-wzzE_Rvr GG333/4LL | CACATAATCATCAGGAAGG |
| pZS21-wzzE_fwd G339A | GATCGGGGCTgctGTCGCATTAACCC |
| pZS21-wzzE_Rvr G339A | AGCCCCCCGACAATGCCC |
| pwzzE_x3F Fwd | TGATAAGTGGATCCGTCGA |
| pwzzE_x3F Rvr | ATTGGGCGGCAACGAGCTTT |
| WzzE: Cmr Fp | TAACCCGCCGTTGCTCGAAAATGGAGAAAAAAATCACTGG |
| WzzE: Cmr Rp | CCACTCCCACCGCTGCCTCCCGCCCCGCCCTGCCACTCAT |
| Cmr: GS-x3F FP | GGAGGCAGCGGTGGGAGTGGCGGAGACTACAAGGACCACGA |
